# Visual experience-driven switch in endothelial metabolic transporters is essential for cerebrovascular maturation

**DOI:** 10.64898/2026.10.26.762059

**Authors:** Namsuk Kim, Eun-Young Choi, Sehyun Chae, Jin-Young Jeong, Hyong Kyu Kim, Won-Jong Oh

**Author notes:** These authors contributed equally to this work (Namsuk Kim, Eun-Young Choi, Sehyun Chae). Corresponding author: Namsuk Kim, Neurovascular Biology Laboratory, Department of Neuroscience, Korea Brain Research Institute, Daegu 41062, South Korea.; Won-Jong Oh, Neurovascular Biology Laboratory, Department of Neuroscience, Korea Brain Research Institute, Daegu 41062, South Korea.

## Abstract

Angiogenesis and blood‒brain barrier (BBB) maturation are tightly regulated by coordinated interactions among neuronal, glial, and endothelial cells. Although neuronal activity is essential for postnatal cerebrovascular development, how it is translated into endothelial programs that coordinate vascular maturation remains unknown. Here, we show that visual experience drives metabolic reprogramming in brain endothelial cells by decreasing the lactate transporter MCT1 while increasing the glucose transporter GLUT1, a switch that is essential for proper vascular development. Sustained endothelial expression of MCT1 impaired vascular development and BBB maturation, resulting in vascular leakage. Mechanistically, the developmental decline in TGF-β signaling drives the endothelial MCT1-to-GLUT1 transition. Endothelial-derived TGF-β1 further reinforces the immature angiogenic state through a lactate-dependent autocrine positive feedback loop, whereas sensory experience promotes astrocyte maturation, which attenuates endothelial TGF-β signaling, potentially by changing perivascular environment. Together, our findings identify a neuron‒astrocyte‒endothelial metabolic signaling axis that links sensory experience to endothelial metabolic reprogramming and coordinates the transition from angiogenesis to BBB maturation.

## Introduction

The formation of a functional cerebrovascular network is a hallmark of brain development, characterized by a highly orchestrated balance between angiogenesis and the maturation of the blood‒brain barrier (BBB)^1‒6^. This process does not occur in isolation but is tightly regulated by coordinated interactions among neurons, glia, and endothelial cells. Recent spatial profiling of the human brain vasculature further revealed that vascular cells are organized into specialized cellular ensembles, highlighting the spatially coordinated organization of the neurovascular environment^7^. Emerging evidence suggests that sensory-driven neuronal activity plays an important role in shaping this vascular landscape during postnatal development^4,8^. For example, eye opening around postnatal day 14 (P14) in mice triggers a surge of neuronal activity in the visual cortex that coincides with a critical period of cerebrovascular remodeling^9,10^. However, despite growing evidence linking neuronal activity to cerebrovascular development, the molecular mechanisms by which sensory experience directs the transition from an angiogenic state to a mature, restrictive BBB remain largely unknown.

Among the metabolic substrates delivered through the BBB, glucose and lactate play central roles in supporting brain function. Whereas glucose serves as the predominant energy source for the mature brain, lactate functions both as an alternative metabolic substrate during development and as an important signaling molecule in physiological and pathological contexts^11‒15^. Their transport across the BBB is mediated primarily by the endothelial transporters, Glucose Transporter type I (GLUT1) and Monocarboxylate Transporter type 1 (MCT1), respectively. During brain development, brain endothelial cells undergo a characteristic transition from an MCT1-high to a GLUT1-high transporter profile, coinciding with the developmental shift in cerebral fuel utilization from lactate and ketone bodies toward glucose^16,17^. As in many developing tissues, this metabolic transition is likely to represent an actively regulated developmental program rather than a passive adaptation to changing substrate availability^18,19^. However, the molecular signals that initiate and coordinate this endothelial metabolic switch remain largely unknown because of the complex reciprocal communication among multiple cell types within the neurovascular unit (NVU).

Transforming growth factor-β (TGF-β) signaling is a key regulator of multiple aspects of cerebrovascular biology, including endothelial proliferation, differentiation, and junctional organization. Dysregulation of this pathway can lead to vascular instability and diverse cerebrovascular disorders ^20‒22^. Although the roles of TGF-β signaling in embryonic cerebrovascular development have been extensively investigated, its functions in postnatal cerebrovascular development and maturation remain relatively understudied^6,23^. Beyond its vascular functions, TGF-β signaling also regulates cellular metabolism, including glucose and lipid metabolism, although its effects differ across cell types^24‒26^. Previous studies have shown that TGF-β can regulate glucose transporter expression, glucose uptake, and glycolytic activity, as well as pathways involved in lipid synthesis and fatty acid oxidation, although these effects vary depending on cellular context^24‒26^. Together, these observations suggest a potential intersection between TGF-β signaling, metabolic regulation, and cerebrovascular development, although how these processes are integrated during brain development remains poorly understood.

Here, we show that sensory experience‒driven neuronal activity induces an endothelial MCT1-to-GLUT1 switch required for normal angiogenesis and BBB maturation. Mechanistically, endothelial TGF-β1 sustains a transient MCT1-high angiogenic state through a lactate-dependent positive-feedback loop. Sensory experience promotes astrocyte maturation and MCT1 expression, potentially reducing perivascular lactate availability and attenuating this loop to enable the endothelial metabolic switch. Together, our findings identify a neuron‒astrocyte‒endothelial signaling axis that links neuronal activity to endothelial metabolic remodeling and coordinates the transition from angiogenesis to BBB maturation.

## Results

### Sensory experience regulates endothelial metabolic programs during postnatal vascular development

To investigate the underlying mechanism by which neural activation influences the brain endothelium, we examined gene expression profiles during visual cortex development as a model of natural neural activation. In mice, eye opening typically occurs around postnatal day 14 (P14), and the subsequent exposure to light triggers neural activation in the visual cortex^9,10^. Thus, we performed translatome analysis on the brain endothelial cells (BECs) from the visual cortex at postnatal day 10 (P10), prior to eye opening, and at postnatal day 30 (P30), after eye opening, using an optimized RiboTag-based approach (Figure 1A and Figure S1A)^27^. The enrichment and specificity of BECs in the isolated mRNA pool were validated by reverse transcription polymerase chain reaction (RT-PCR) and quantitative reverse transcription PCR (RT-qPCR) for endothelial and non-endothelial cell markers (Figures S1B and S1C).

**Figure 1.**
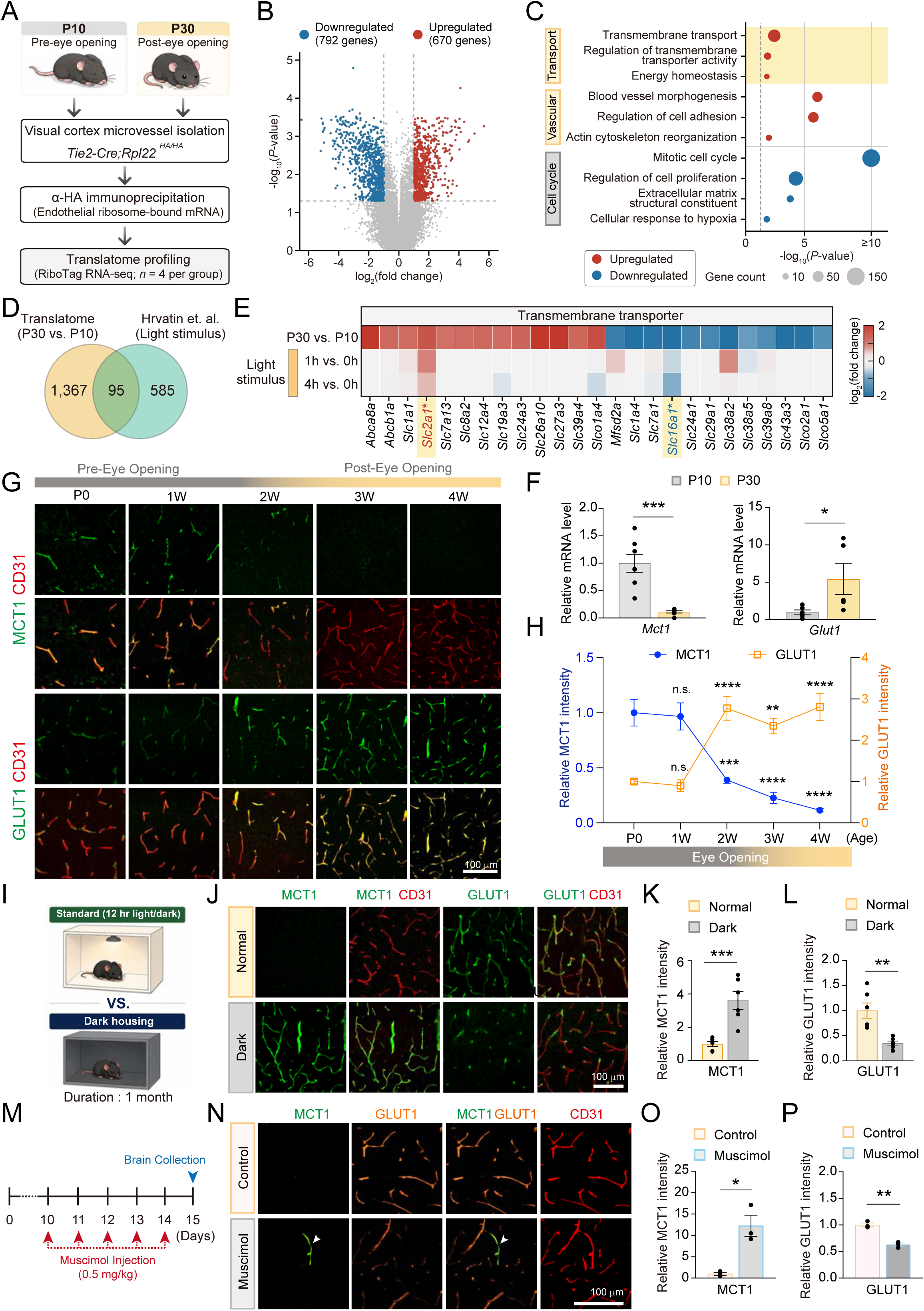
Sensory experience regulates a reciprocal lactate-to-glucose transporter switch in brain endothelium. (A) Schematic of RiboTag-based translatome profiling of brain endothelial cells. RiboTag-based endothelial translatome was isolated from visual cortex microvessels of *Tie2-Cre;Rpl22^HA/HA^* mice at P10 (pre-eye opening) and P30 (post-eye opening) by anti-HA immunoprecipitation and sequenced (n = 4 mice per group). (B) Volcano plot of the brain endothelial translatome at P30 relative to P10. Genes upregulated and downregulated at P30 are shown in red and blue, respectively; genes that were not differentially translated are shown in gray. Dashed lines indicate the fold change and significance thresholds. The number of genes in each group is indicated. (C) Functional over-representation analysis of the up- and downregulated genes. Dot color indicates up-(red) or downregulation (blue) at P30. Dot size indicates the number of genes annotated to each term, and the x axis indicates enrichment significance (values above 10 are shown as ≥10). The dashed line indicates *P* = 0.05. (D) Overlap between genes differentially translated in brain endothelial cells between P30 and P10 and light-responsive endothelial genes identified from single-cell RNA sequencing of the light-stimulated visual cortex^29^. The number of genes in each group is indicated. (E) Heat map showing transmembrane transporters among the genes differentially translated between P30 and P10. Rows show the developmental change in the endothelial translatome (P30 versus P10) and the responses to light stimulation in brain endothelial cells (1 h and 4 h versus 0 h). Color indicates log2-fold change, with red and blue denoting up- and downregulation, respectively. Asterisks indicate transporters that were significantly regulated in both datasets with a concordant expression pattern. (F) RT-qPCR analysis of *Mct1* and *Glut1* in RiboTag-isolated endothelial RNA from the visual cortex of P10 and P30 mice. For *Mct1*, n = 7 mice per group; for *Glut1*, n = 6 mice at P10 and n = 5 mice at P30. Unpaired two-tailed Student’s *t* test. (G) Representative immunofluorescence images of MCT1 or GLUT1 together with the endothelial marker CD31 in the visual cortex at the indicated postnatal ages. The approximate timing of eye opening is indicated above the images. Scale bar, 100 μm. (H) Quantification of relative MCT1 and GLUT1 fluorescence intensity in CD31-positive blood vessels at the indicated postnatal ages. The approximate timing of eye opening is indicated below the graph. n = 3 mice per time point, with two sections analyzed per mouse. One-way ANOVA with Tukey’s multiple-comparisons test. (I) Experimental design for sensory deprivation. Mice were maintained under standard 12-h light/12-h dark conditions (Normal) or dark-rearing conditions (Dark) for 1 month. (J) Representative immunofluorescence images of MCT1, GLUT1, and CD31 in the visual cortex of normally reared and dark-reared mice. Scale bar, 100 μm. (K) and L) Quantification of relative endothelial MCT1 (K) and GLUT1 (L) levels in normally reared and dark-reared mice. For MCT1, n = 6 mice; for GLUT1, n = 8 mice, with two sections analyzed per mouse. Unpaired two-tailed Student’s *t* test. (M) Experimental design for pharmacological suppression of neuronal activity. Mice received daily intraperitoneal injections of muscimol, a GABA_A_ receptor agonist, (0.5 mg/kg) from P10 to P14, followed by brain collection at P15. (N) Representative immunofluorescence images of MCT1, GLUT1, and CD31 in the visual cortex of control and muscimol-treated mice. Arrowheads indicate MCT1-positive blood vessels in muscimol-treated mice. Scale bar, 100 μm. (O and P) Quantification of relative endothelial MCT1 (O) and GLUT1 (P) levels in CD31-positive blood vessels from control and muscimol-treated mice. For both MCT1 and GLUT1, n = 3 mice per group. Unpaired two-tailed Student’s *t* test. Data are presented as mean ± SEM; \**p* < 0.05, \*\**p* < 0.01, \*\*\**p* < 0.001, \*\*\*\**p* < 0.0001 and n.s., not significant. See also Figures S1 and S2.

Principal component analysis of the endothelial translatome profiles clearly separated the P10 and P30 stages (Figure S1D). A total of 1,462 genes were differentially translated between the two stages, with 670 upregulated and 792 downregulated at P30 (Figure 1B; Table S1), indicating extensive remodeling of the endothelial gene expression program. To identify the cellular processes represented among these genes, we performed over-representation analysis of the up- and downregulated genes using ConsensusPathDB^28^. The downregulated genes were enriched for terms related to the mitotic cell cycle, cell proliferation, extracellular matrix composition, and the cellular response to hypoxia. In contrast, the upregulated genes were enriched for processes related to blood vessel morphogenesis, cell adhesion, and actin cytoskeleton reorganization, together with transport-related processes including transmembrane transport, regulation of transmembrane transporter activity, and energy homeostasis (Figure 1C).

To identify endothelial genes responsive to neural activation, we next reanalyzed a published single-cell RNA sequencing dataset of the visual cortex, in which dark-adapted mice were re-exposed to light to evoke neural activation^29^. Analysis of the endothelial population revealed genes whose expression changed 1 and 4 h after light exposure (Figures S1E and S1F). By comparing the two datasets, we identified 95 genes that were both differentially translated between P30 and P10 and responsive to light stimulation, representing candidate endothelial programs regulated by neural activation (Figure 1D). Because transport-related processes were prominently enriched in the translatome, we next examined whether the differentially translated transporters changed in the single-cell dataset. Only two transporters changed in both datasets: *Slc16a1*, encoding the lactate transporter MCT1, decreased from P10 to P30 and after light exposure, whereas *Slc2a1*, encoding the glucose transporter GLUT1, showed the opposite pattern (Figure 1E; Figure S1G). We therefore focused on MCT1 and GLUT1, which transport the principal energy substrates of the developing brain. To validate the developmental expression changes identified by the translatome analysis, we performed RT-qPCR analysis of nine candidate genes using RiboTag-isolated endothelial RNA from the P10 and P30 visual cortices. In addition to *Mct1* and *Glut1*, *Hspb1*, *Timp3*, *Mal*, and *Rpl38* also showed significant changes in expression between P10 and P30(Figure 1F; Figures S2A and S2B).

We next examined the developmental expression of MCT1 and GLUT1 at the protein level. Immunofluorescence analysis of the visual cortex across postnatal development (P0, 1, 2, 3, and 4 weeks) showed that endothelial MCT1 expression was high during the early postnatal period and declined markedly around eye opening, whereas endothelial GLUT1 expression increased reciprocally (Figures. 1G and 1H). This pattern is consistent with previous reports of the developmental regulation of these transporters during postnatal brain maturation^17^. In contrast, the total protein levels of both transporters progressively increased in whole visual cortex lysates, suggesting that the reciprocal transporter switch is specific to brain endothelial cells (Figures S2C and S2D).

To determine whether visual experience contributes to the postnatal switch in transporter expression, we analyzed MCT1 and GLUT1 expression in the visual cortex of dark-reared mice at P30 (Figure 1I). Under dark-rearing conditions, c-Fos expression, a marker of neuronal activation, was markedly reduced, confirming effective suppression of visual experience‒evoked neuronal activity^30^ (Figures S2E and S2F). Consistent with this, the developmental decline in MCT1 expression was attenuated, whereas the developmental increase in GLUT1 expression was significantly reduced compared with normally reared mice (Figures 1J‒L). To further determine whether neuronal activity *per se* regulates this transporter switch, we inhibited neuronal activity around the time of eye opening by intraperitoneal administration of muscimol, a GABA_A_ receptor agonist (Figure 1M). Muscimol treatment markedly reduced c-Fos expression (Figures S2G and S2H), confirming effective suppression of neuronal activity, and partially preserved endothelial MCT1 expression in a subset of vessels while attenuating the developmental upregulation of GLUT1 (Figures 1N‒P). In particular, vessels retaining MCT1 expression exhibited markedly lower GLUT1 expression, further supporting a reciprocal relationship between the two transporters. Together, these results support the notion that neuronal activity elicited by visual experience contributes to the reciprocal postnatal regulation of MCT1 and GLUT1 in brain endothelial cells.

### Sustained endothelial MCT1 expression impairs cerebrovascular maturation

Next, we asked whether the developmental downregulation of MCT1 is required for normal cerebrovascular maturation. To address this question, we disrupted the physiological developmental decline in MCT1 by sustaining its expression specifically in brain endothelial cells through postnatal week 2 using the CNS endothelial cell-targeting AAV-BI30 vector^31^ (Figure 2A). Sustained endothelial MCT1 overexpression markedly reduced vascular density, vessel diameter, and vessel length, accompanied by decreased expression of the pan-endothelial marker CD31 compared with AAV-GFP controls (Figures 2B‒E, Figures S3A and S3B).

**Figure 2.**
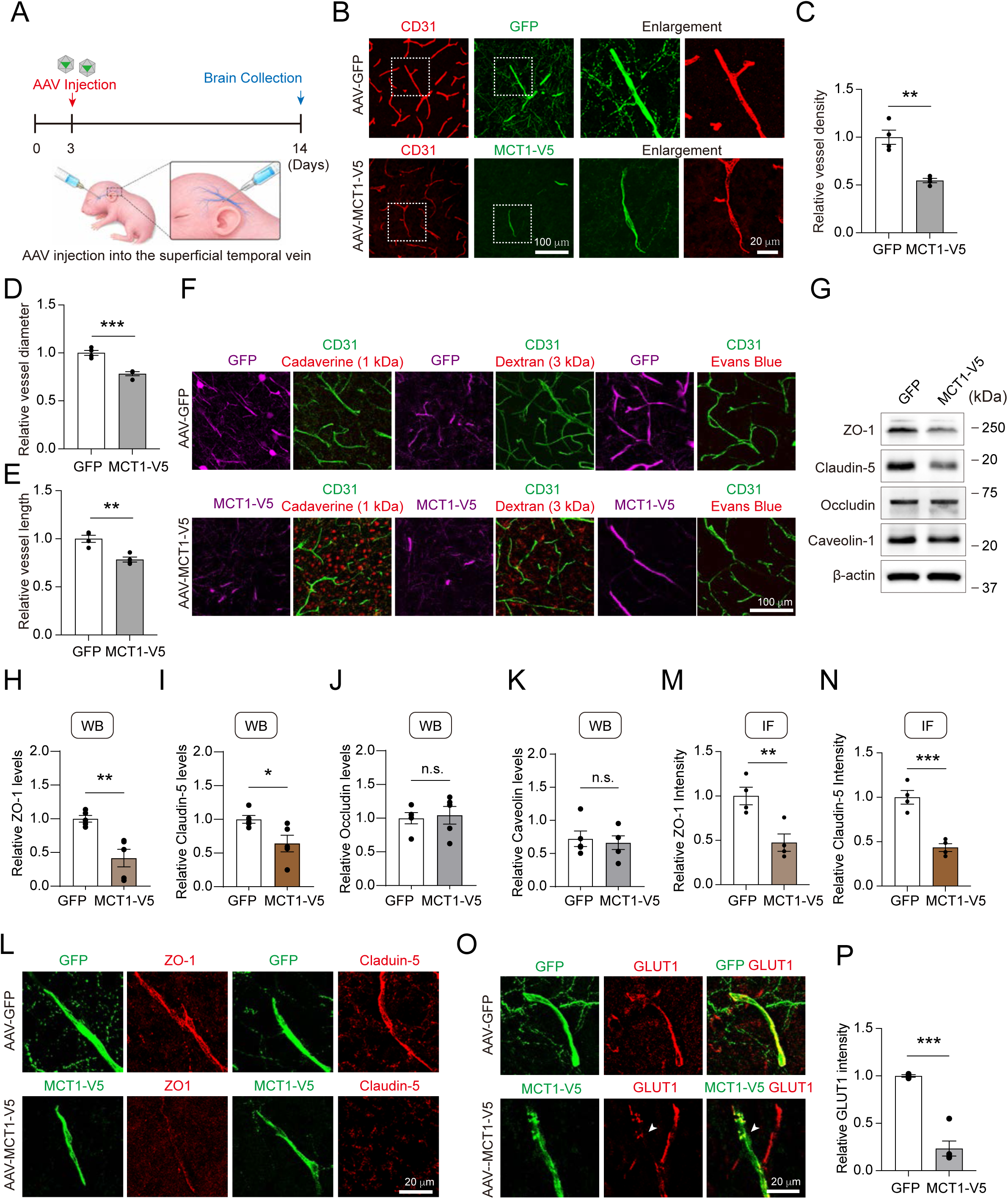
Sustained endothelial MCT1 expression compromises cerebrovascular development. (A) Experimental scheme for sustained MCT1 expression in brain endothelial cells. AAV-GFP or AAV-MCT1-V5 was administered via the superficial temporal vein at P3, followed by brain collection at P14. (B) Representative immunofluorescence images of CD31 and GFP or V5-tagged MCT1 in the visual cortex of AAV-GFP-and AAV-MCT1-V5-injected mice. Boxed regions are shown at higher magnification. Scale bars, 100 μm (overview) and 20 μm (enlargement). (C‒E) Quantification of vessel density (C), vessel diameter (D), and vessel length (E) in AAV-GFP- and AAV-MCT1-V5-injected mice. n = 4 mice per group, with ten sections analyzed and averaged per mouse. Each mouse was considered an independent biological replicate. Unpaired two-tailed Student’s *t* test. (F) Representative images of BBB permeability assessed using cadaverine (1 kDa), dextran (3 kDa), and Evans blue in the visual cortex of AAV-GFP- and AAV-MCT1-V5-injected mice. Similar findings were observed in three mice per group. Scale bar, 100 μm. (G) Representative western blots of ZO-1, Claudin-5, Occludin, and Caveolin-1 in brain lysates from AAV-GFP- and AAV-MCT1-V5-injected mice. (H‒K) Quantification of ZO-1 (H), Claudin-5 (I), Occludin (J), and Caveolin-1 (K) protein levels normalized to β-actin. n = 5 mice per group. Unpaired two-tailed Student’s *t* test. (L) Representative immunofluorescence images of ZO-1 and Claudin-5 in GFP- or MCT1-V5-positive blood vessels in the visual cortex of AAV-GFP- and AAV-MCT1-V5-injected mice, respectively. Scale bar, 20 μm. (M and N) Quantification of relative ZO-1 (M) and Claudin-5 (N) fluorescence intensity in GFP- or MCT1-V5-positive blood vessels. n = 4 mice per group, with two sections analyzed per mouse. Unpaired two-tailed Student’s *t* test. (O) Representative immunofluorescence images of GLUT1 in GFP- or MCT1-V5-positive blood vessels at P14 after AAV injection. Scale bar, 20 μm. (P) Quantification of relative GLUT1 fluorescence intensity in GFP- or MCT1-V5-positive blood vessels. n = 3 mice for GFP and 5 mice for MCT1-V5, with two sections analyzed per mouse. Unpaired two-tailed Student’s *t* test. Data are presented as mean ± SEM; \**p* < 0.05, \*\**p* < 0.01, \*\*\**p* < 0.001, and n.s., not significant. See also Figure S3.

Moreover, sustained endothelial MCT1 overexpression increased the extravasation of low-molecular-weight tracers, including Alexa Fluor 555‒conjugated cadaverine (∼1 kDa) and Texas Red‒conjugated dextran (3 kDa), whereas no leakage of endogenous immunoglobulin G (IgG) or Evans Blue was detected (Figure 2F and Figure S3C). Consistent with these findings, the expression of the tight junction proteins, zonula occludens-1 (ZO-1) and Claudin-5, was significantly reduced, as confirmed by both western blotting and immunofluorescence (Figures 2G‒N). In line with the effects of muscimol treatment (Figures 1N‒P), GLUT1 expression was also dramatically suppressed in endothelial cells overexpressing MCT1 (Figures 2O and 2P). These findings suggest that sustained endothelial MCT1 expression disrupts the normal developmental regulation of GLUT1.

Given the BBB defects observed following sustained endothelial MCT1 expression, we next investigated whether a similar phenotype occurs under dark-rearing conditions, which also prevent the developmental downregulation of endothelial MCT1 (Figure 1). Indeed, dark-reared mice exhibited increased extravasation of fluorescent tracers into the brain parenchyma compared with normally reared controls (Figure S3D). Together, these findings demonstrate that the developmental downregulation of endothelial MCT1 is essential for postnatal endothelial metabolic reprogramming and proper cerebrovascular maturation, including the establishment of BBB integrity.

### Endothelial TGF-β signaling orchestrates the reciprocal transition in MCT1 and GLUT1 expression

To identify the upstream regulator coordinating the opposing expression of these transporters in response to visual experience‒driven neuronal activation, we screened a panel of factors known to participate in vascular development and angiogenesis for their ability to regulate MCT1 and GLUT1 in cultured brain endothelial cells. hCMEC/D3 cells were treated with TGF-β1, IGF-I, IGF-II, VEGF, BDNF, or LiCl (lithium chloride; WNT agonist) and MCT1 and GLUT1 protein levels were assessed by western blotting (Figure S4). Among the factors tested, TGF-β1 induced the most prominent increase in MCT1 expression, whereas IGF-I and IGF-II caused smaller increases. LiCl modestly but significantly increased both MCT1 and GLUT1 levels, whereas VEGF and BDNF had little effect (Figures S4B‒S4F). In contrast, GLUT1 expression did not change consistently in response to any of the factors, including TGF-β1 (Figures S4C and S4F). We reasoned that the absence of further GLUT1 downregulation might reflect the fact that endogenous TGF-β signaling already suppresses GLUT1 expression under basal culture conditions, such that further stimulation with exogenous TGF-β1 would not reduce it further. If so, blocking TGF-β signaling should relieve this suppression and reveal reciprocal regulation of the two transporters. To test this possibility, we treated hCMEC/D3 cells with SIS3, a SMAD3 inhibitor, or galunisertib, a TGF-β receptor type I inhibitor. Both treatments markedly reduced MCT1 expression while increasing GLUT1 expression (Figures 3A‒F), indicating that MCT1 and GLUT1 are oppositely regulated by canonical TGF-β signaling.

**Figure 3.**
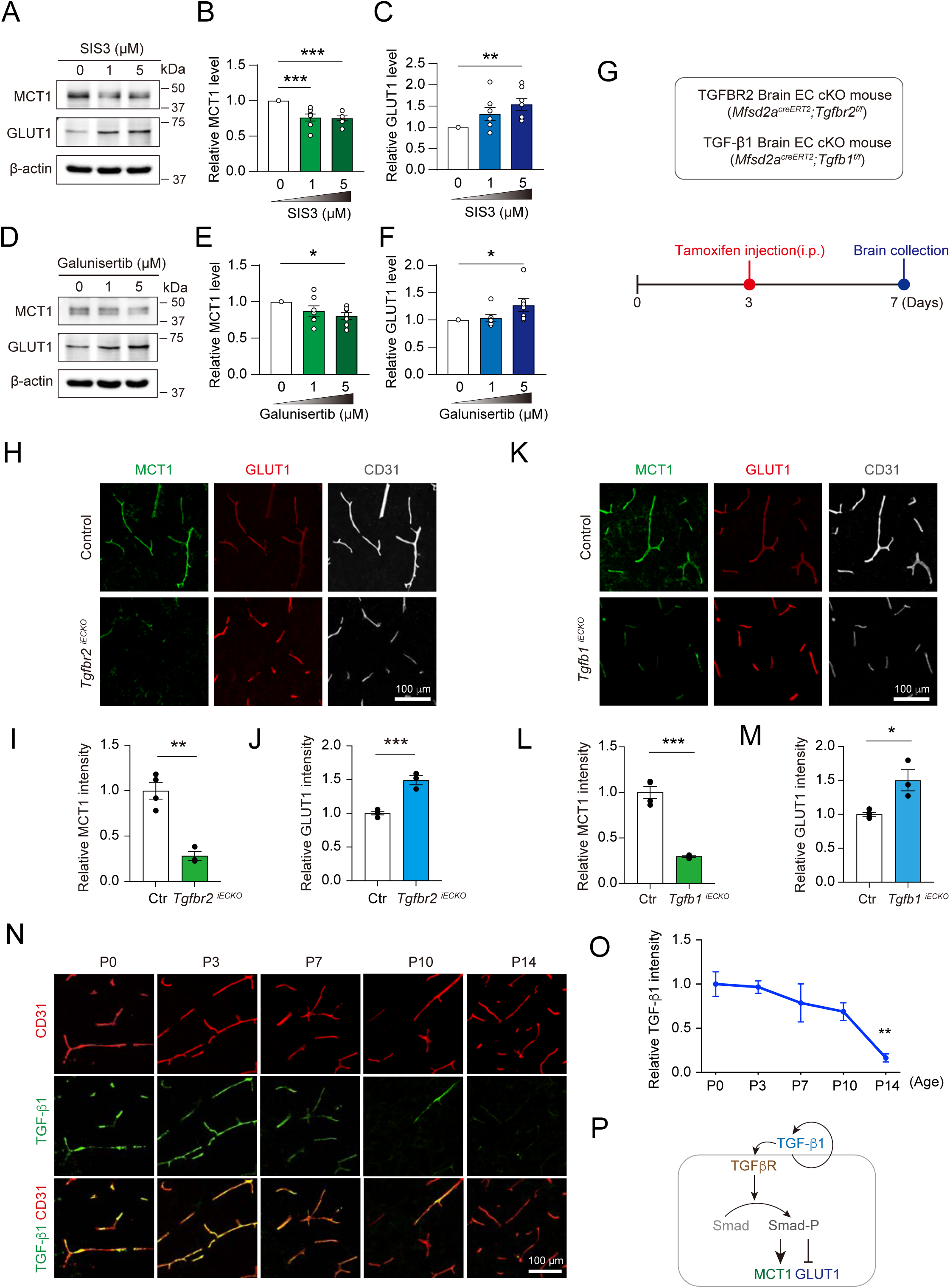
Endothelial-derived TGF-β1 regulates the MCT1-to-GLUT1 switch in brain endothelial cells. (A) Representative western blots of MCT1 and GLUT1 in hCMEC/D3 cells treated with the SMAD3 inhibitor SIS3 at the indicated concentrations. β-actin was used as a loading control. (B and C) Quantification of MCT1 (B) and GLUT1 (C) protein levels following SIS3 treatment, normalized to β-actin. n =7 biological samples per group. (D) Representative western blots of MCT1 and GLUT1 in hCMEC/D3 cells treated with the TGF-β receptor I inhibitor galunisertib at the indicated concentrations. β-actin was used as a loading control. (E and F) Quantification of MCT1 (E) and GLUT1 (F) protein levels following galunisertib treatment, normalized to β-actin. n = 7 biological samples per group. (G) Experimental scheme for inducible endothelial-specific deletion of *Tgfbr2* or *Tgfb1*. Tamoxifen was administered intraperitoneally at P3, and brains were collected at P7. (H) Representative immunofluorescence images of MCT1, GLUT1, and CD31 in the visual cortex of control and endothelial-specific *Tgfbr2* conditional knockout (*Tgfbr2* ^1ECKO^) mice. Scale bar, 100 μm. (I) and J) Quantification of relative endothelial MCT1 (I) and GLUT1 (J) fluorescence intensity in control and *Tgfbr2* ^iECKO^ mice. n = 4 mice for control and n = 3 mice for *Tgfbr2* ^iECKO^, with two sections analyzed per mouse. Unpaired two-tailed Student’s *t* test. (K) Representative immunofluorescence images of MCT1, GLUT1, and CD31 in the visual cortex of control and endothelial-specific *Tgfbl* conditional knockout (*Tgfbl* ^1ECKO^) mice. Scale bar, 100 μm. (L) and M) Quantification of relative endothelial MCT1 (L) and GLUT1 (M) fluorescence intensity in control and *Tgfb1* ^iECKO^ mice. n = 4 mice per group, with two sections analyzed per mouse. Unpaired two-tailed Student’s *t* test. (N) Representative immunofluorescence images of TGF-β1 and CD31 in the cerebral cortex at P0, P3, P7, P10, and P14. Scale bar, 100 μm. (O) Quantification of relative endothelial TGF-β1 fluorescence intensity at the indicated postnatal ages. n = 4 mice per age group, with two sections analyzed per mouse. One-way ANOVA followed by Dunnett’s multiple-comparisons test. (P) Summary model of autocrine TGF-β signaling that upregulates MCT1 and downregulates GLUT1 expression in brain endothelial cells. Data are presented as mean ± SEM; \**p* < 0.05, \*\**p* < 0.01, and \*\*\**p* < 0.001. See also Figures S4 and S5.

To test whether this reciprocal regulation also occurs *in vivo*, we generated a brain endothelial cell-specific TGF-β receptor type II (*Tgfbr2*) knockout mouse model^32^. To avoid lethality, *Tgfbr2^flox/flox^* mice were crossed with *Mfsd2a-CreERT2* mice, in which tamoxifen-inducible Cre recombinase is predominantly expressed in BECs^33,34^. Because upregulation of MCT1 and suppression of GLUT1 were observed during early postnatal development before eye opening, tamoxifen was administered at P3, and immunostaining was performed at P7, a developmental stage characterized by high MCT1 expression (Figure 3G). Immunostaining confirmed efficient loss of TGFBR2 expression in brain endothelial cells following tamoxifen-induced deletion (Figure S5B). In *Tgfbr2*-deficient neonates, MCT1 expression in brain endothelial cells was markedly reduced, whereas GLUT1 levels were slightly upregulated, indicating that TGF-β signaling is required to maintain the balance toward an MCT1-high/GLUT1-low state in brain endothelial cells during early development. (Figures 3H‒J).

Given the opposing effects of TGF-β signaling on endothelial MCT1 and GLUT1 expression, we next asked whether endogenous endothelial TGF-β signaling undergoes developmental changes that parallel the reciprocal transition in MCT1 and GLUT1 expression during postnatal cerebrovascular maturation. To address this, we examined phosphorylated SMAD2 (p-SMAD2), a key downstream readout of canonical TGF-β signaling^35^, in the visual cortex from birth (P0) through 4 weeks of age. Endothelial p-SMAD2 levels were high before eye opening but declined markedly following eye opening and the onset of visual experience‒evoked neuronal activity (Figures S5C and S5D). This developmental decline closely paralleled the downregulation of endothelial MCT1 and reciprocal upregulation of GLUT1, supporting a link between attenuation of TGF-β signaling and the endothelial MCT1-to-GLUT1 transition during postnatal cerebrovascular maturation. Together with the pharmacological inhibition and endothelial *Tgfbr2* knockout results, these findings support a model in which developmental attenuation of endothelial TGF-β signaling contributes to the reciprocal transition from an MCT1-high/GLUT1-low to an MCT1-low/GLUT1-high state during postnatal cerebrovascular maturation. We next sought to determine which cell type provides the TGF-β ligands during early postnatal development.

During brain development, the sources of TGF-β1 ligands exhibit marked spatiotemporal diversity, originating from multiple cell types, including radial glial cells (RGCs), microglia, and astrocytes-depending on the developmental stage and spatial context^36‒38^. However, the cellular source of TGF-β1 responsible for regulating endothelial MCT1 and GLUT1 expression during early postnatal cerebrovascular maturation remains unclear. To identify the cellular source of TGF-β1 that regulates brain endothelial cells during the early postnatal period, we generated cell type-specific conditional knockouts of *Tgfb1* by crossing floxed *Tgfb1* mice with various Cre driver lines — *Mfsd2a-CreERT2* (endothelial cells), *Pdgfrb-CreERT2* (pericytes), *Aldh1l1-CreERT2* (astrocytes), and *Camk2a-CreERT2* (neurons). Among these models, endothelial-specific deletion of *Tgfb1* markedly reduced endothelial MCT1 expression (Figures 3K-M). In contrast, astrocyte-specific deletion of *Tgfb1* caused only a modest reduction in endothelial MCT1 expression, whereas pericyte- or neuron-specific *Tgfb1* deletion had no detectable effect (Figures S5E‒I). These findings identify endothelial-derived TGF-β1 as the predominant functional source maintaining the MCT1-high/GLUT1-low endothelial state during early postnatal development, with astrocyte-derived TGF-β1 making a comparatively minor contribution. Consistent with this functional evidence, immunostaining at P7 revealed prominent TGF-β1 expression in brain endothelial cells, which was markedly diminished in *Mfsd2a-CreERT2;Tgfb1^fl/fl^* mice, confirming efficient endothelial deletion and further supporting endothelial cells as the predominant source of TGF-β1 during early postnatal development (Figure S5J). Moreover, endothelial TGF-β1 expression was high during early postnatal development but declined markedly by P14 (Figures 3N and 3O), consistent with the developmental attenuation of endothelial p-SMAD2 signaling and the reciprocal transition in MCT1 and GLUT1 expression (Figures S5C and S5D). Collectively, these findings establish TGF-β signaling as a key developmental regulator of the reciprocal MCT1/GLUT1 transition in brain endothelial cells and identify endothelial-derived TGF-β1 as the predominant ligand source sustaining MCT1 expression during early postnatal development (Figure 3P). Consistent with this immature endothelial state, prolonged TGF-β1 treatment also reduced transendothelial electrical resistance and decreased ZO-1 and Claudin-5 expression in hCMEC/D3 cells, whereas Occludin levels were largely unchanged (Figure S6). Together with our findings that sustained endothelial MCT1 overexpression impairs vascular remodeling, reduces tight junction protein expression, and increases BBB permeability to low-molecular-weight tracers (Figures 2B‒N), these results suggest that developmental attenuation of endothelial TGF-β signaling may facilitate cerebrovascular maturation and BBB integrity, at least in part by permitting the MCT1-to-GLUT1 transition.

### Endothelial TGF-β1‒MCT1‒lactate signaling forms a positive feedback loop

Previous studies have demonstrated that TGF-β1 can induce its own expression through autoregulatory mechanisms in multiple cell types^39‒41^. We therefore examined whether a similar mechanism operates in brain endothelial cells using the human brain microvascular endothelial cell line hCMEC/D3. Treatment with active TGF-β1 increased SMAD2/3 phosphorylation, confirming activation of canonical TGF-β signaling, which was accompanied by increased endogenous TGF-β1 expression (Figure S7). To determine whether this autoregulatory mechanism also operates *in vivo*, we examined endothelial cell-specific *Tgfbr2* knockout mice. Consistent with impaired canonical TGF-β signaling, endothelial TGF-β1 expression was markedly reduced at P7 in *Tgfbr2*-deficient mice (Figures 4A‒C). This reduction was accompanied by decreased MCT1 expression (Figures 3H‒J) and lower brain lactate levels (Figure 4D). Given recent evidence that lactate can promote TGF-β1 expression through histone lactylation in other cell types^42,43^, these findings raised the possibility that MCT1-associated lactate accumulation contributes to the maintenance of endothelial TGF-β1 expression.

**Figure 4.**
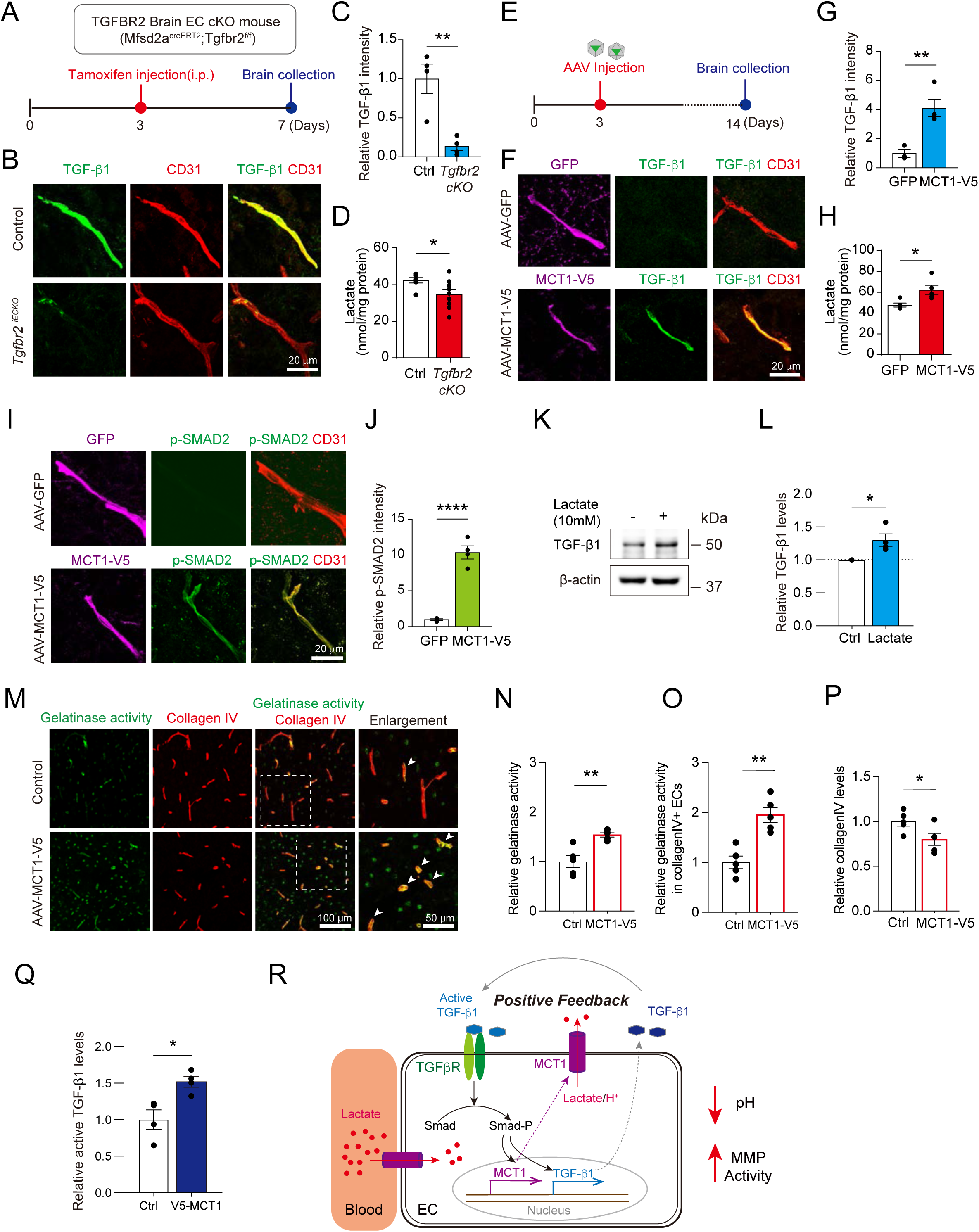
MCT1-Lactate axis activates endothelial TGF-β signaling via perivascular remodeling. (A) Experimental scheme for inducible endothelial deletion of *Tgfbr2*. Tamoxifen was administered intraperitoneally at P3, and brains were collected at P7. (B) Representative immunofluorescence images of TGF-β1 and CD31 in the visual cortex of control and endothelial-specific *Tgfbr2* conditional knockout (*Tgfbr2* ^1ECKO^) mice. Scale bar, 20 μm. (C) Quantification of relative endothelial TGF-β1 fluorescence intensity in control and *Tgfbr2* ^iECKO^ mice. n = 4 mice per group, with two sections analyzed per mouse. Unpaired two-tailed Student’s *t* test. (D) Quantification of brain lactate concentrations, normalized to total protein (nmol/mg protein), in control and *Tgfbr2* ^iECKO^ mice. n = 7 control and n = 9 *Tgfbr2* ^iECKO^ mice. Unpaired two-tailed Student’s *t* test. (E) Experimental scheme for sustained endothelial MCT1 expression. AAV-GFP or AAV-MCT1-V5 was administered at P3, and brains were collected at P14. (F) Representative immunofluorescence images of TGF-β1 in GFP- or MCT1-V5-positive blood vessels in the cerebral cortex of AAV-GFP- and AAV-MCT1-V5-injected mice, respectively. Scale bar, 20 μm. (G) Quantification of relative endothelial TGF-β1 fluorescence intensity in AAV-GFP- and AAV-MCT1-V5-injected mice. n = 3 mice for AAV-GFP and n = 4 mice for AAV-MCT1-V5, with two sections analyzed per mouse. Unpaired two-tailed Student’s *t* test. (H) Quantification of brain lactate concentrations, normalized to total protein (nmol/mg protein), in AAV-GFP- and AAV-MCT1-V5-injected mice. n = 5 mice per group. Unpaired two-tailed Student’s *t* test. (I) Representative immunofluorescence images of phosphorylated SMAD2 (p-SMAD2) in GFP- or MCT1-V5-positive blood vessels in the cerebral cortex of AAV-GFP- and AAV-MCT1-V5-injected mice. Scale bar, 20 μm. (J) Quantification of relative endothelial p-SMAD2 fluorescence intensity in AAV-GFP- and AAV-MCT1-V5-injected mice. n = 4 mice per group, with two sections analyzed per mouse. Unpaired two-tailed Student’s *t* test. (K) Representative western blot of TGF-β1 in the hCMEC/D3 cells treated with 10 mM L-lactate for 48 h. β-actin was used as a loading control. (L) Quantification of TGF-β1 protein levels normalized to β-actin. n = 4 biological samples per group. Unpaired two-tailed Student’s *t* test. (M) Representative images of gelatinase activity and Collagen IV in the cerebral cortex of AAV-GFP- and AAV-MCT1-V5-injected mice. Boxed regions are shown at higher magnification. Arrowheads indicate gelatinase activity associated with blood vessels. Scale bars, 100 μm (overview) and 50 μm (enlargement). (N‒P) Quantification of total gelatinase activity (N), gelatinase activity associated with collagen IV-positive blood vessels (O), and relative collagen IV levels (P) in AAV-GFP- and AAV-MCT1-V5-injected mice. n = 5 mice per group. Unpaired two-tailed Student’s *t* test. (Q) Quantification of relative active TGF-β1 levels in microvessels isolated from the visual cortex of AAV-GFP- or AAV-MCT1-V5-injected mice. n = 4 mice per group. Unpaired two-tailed Student’s *t* test. (R) Schematic model summarizing the proposed relationship among endothelial MCT1, lactate, perivascular remodeling, and TGF-β signaling based on the findings in this figure. Data are presented as mean ± SEM; \**p* < 0.05, \*\**p* < 0.01, and \*\*\*\**p* < 0.0001. See also Figure S7.

To explore this possibility, we examined whether sustained endothelial MCT1 expression enhances TGF-β1 signaling. Endothelial MCT1 overexpression using the AAV-BI30 system through P14 increased endothelial TGF-β1 expression (Figures 4E‒G), brain lactate levels (Figure 4H), and endothelial p-SMAD2 levels (Figures 4I and 4J), despite the developmental decline in canonical TGF-β signaling normally observed at this stage (Figures S5C and S5D). Furthermore, direct lactate treatment increased TGF-β1 protein levels in hCMEC/D3 cells (Figures 4K and 4L), supporting a role for lactate in reinforcing endothelial TGF-β1 signaling. Because MCT1 mediates the coupled transport of lactate and H^+^^44^, we hypothesized that sustained endothelial MCT1 expression might promote perivascular acidification. Previous studies have shown that acidic extracellular conditions can induce MMP9 expression^45^ and that extracellular proteolysis can facilitate latent TGF-β1 activation^41,46^. To explore this possibility, we examined perivascular gelatinase activity and extracellular matrix remodeling. *In situ* gelatinase assays revealed a significant increase in overall gelatinase activity following sustained endothelial MCT1 expression (Figures 4M and 4N), with a more pronounced increase within collagen IV-positive vascular structures (Figures 4M and 4O). This was accompanied by a reduction in collagen IV levels (Figures 4M and 4P). Consistent with these changes, ELISA analysis revealed significantly increased levels of active TGF-β1 following sustained endothelial MCT1 expression (Figure 4Q). Together, these findings are consistent with a model in which MCTl-associated lactate/H^+^ transport promotes perivascular proteolytic remodeling, potentially facilitating latent TGF-β1 activation and reinforcing the autoregulatory feedback loop (Figure 4R).

### Endothelial TGF-β‒MCT1‒lactate signaling promotes early postnatal angiogenesis

Having identified an endothelial TGF-β1‒MCT1‒lactate autoregulatory positive feedback loop, we next investigated its functional significance during early postnatal cerebrovascular development. To this end, we examined endothelial cell-specific *Mct1* and *Tgfbr2* conditional knockout mice, in which tamoxifen was administered at P3 and cerebrovascular morphology was analyzed at P7 (Figure 5A). Endothelial-specific deletion of *Mct1* significantly reduced vessel diameter, vascular density, and vessel length compared with control littermates (Figures 5B‒E). Similarly, endothelial-specific deletion of *Tgfbr2* resulted in significant reductions in all three vascular parameters (Figures 5F‒I). These findings indicate that both endothelial MCT1 and TGF-β signaling are required for normal cerebrovascular growth during early postnatal development. To further investigate whether MCT1-mediated lactate transport contributes to the angiogenic properties of brain endothelial cells, we performed scratch-wound migration and tube formation assays. Treatment of hCMEC/D3 cells with 10 mM lactate significantly enhanced cell migration and tube formation, whereas co-treatment with AZD3965, an MCT1 inhibitor, markedly attenuated these responses (Figures S8A‒F). Lactate treatment also increased HIF-1α expression, consistent with previous findings linking lactate signaling to angiogenesis^47^, and upregulated MCT1 expression, without further suppressing GLUT1 expression (Fig. S8G and S8H). These results further support a role for MCT1-mediated lactate transport in promoting the angiogenic properties of brain endothelial cells.

**Figure 5.**
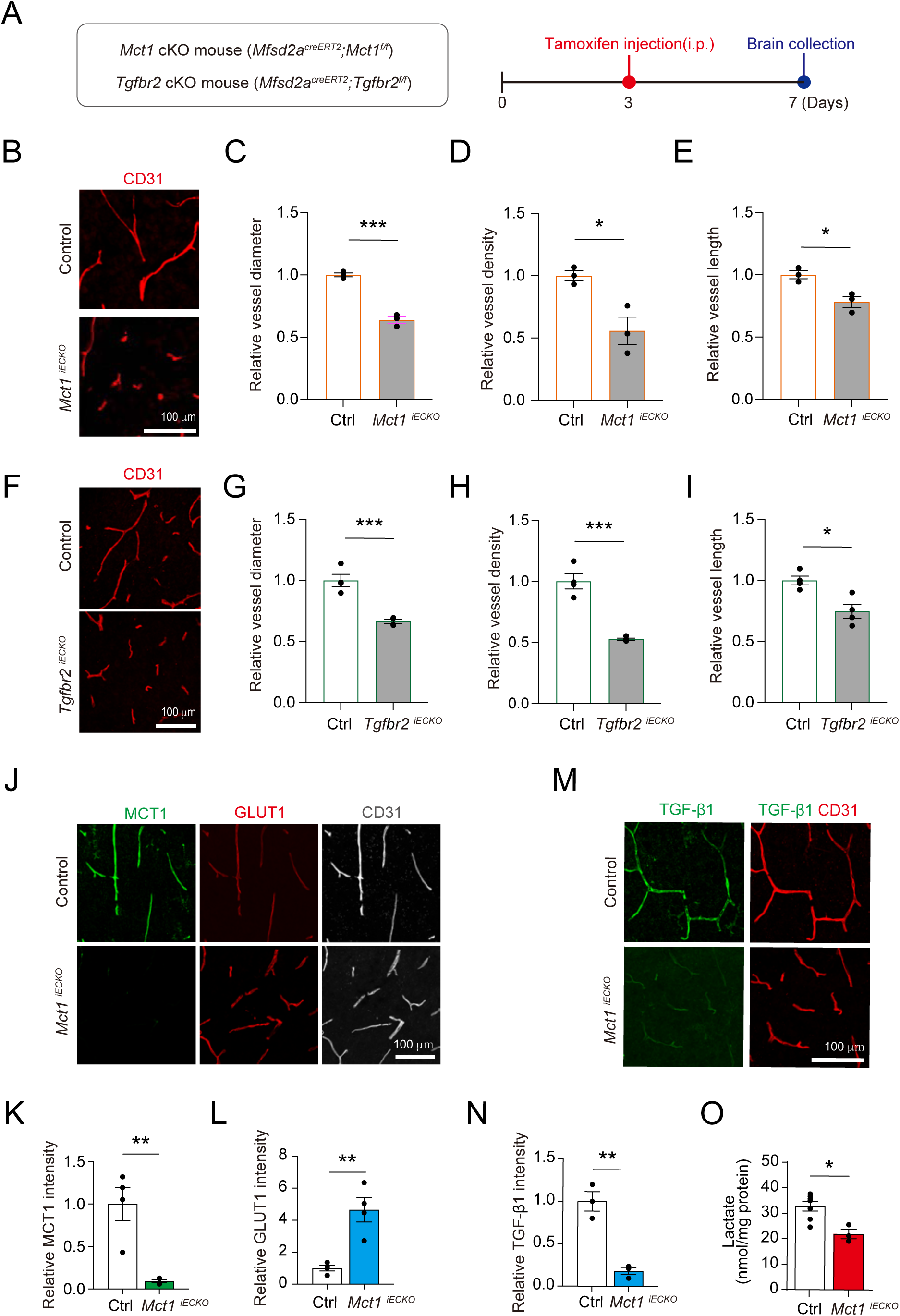
Endothelial TGF-β1‒MCT1‒lactate signaling forms a positive feedback loop. (A) Experimental scheme for inducible endothelial-specific deletion of *Mct1* or *Tgfbr2*. Tamoxifen was administered intraperitoneally at P3, and brains were collected at P7. (B) Representative immunofluorescence images of CD31 in the visual cortex of control and endothelial-specific *Mctl* conditional knockout (*Mctl*^1ECKO^) mice. Scale bar, 100 μm. (C‒E) Quantification of vessel diameter (C), vessel density (D), and vessel length (E) in control and Mct1^iECKO^ mice. n = 3 mice per group, with ten sections analyzed and averaged per mouse. Each mouse was considered an independent biological replicate. Unpaired two-tailed Student’s *t* test. (F) Representative immunofluorescence images of CD31 in the visual cortex of control and endothelial-specific *Tgfbr2* conditional knockout (*Tgfbr2* ^1ECKO^) mice. Scale bar, 100 μm. (G‒I) Quantification of vessel diameter (G), vessel density (H), and vessel length (I) in control and *Tgfbr2* ^iECKO^ mice. n = 4 mice per group, with ten sections analyzed and averaged per mouse. Each mouse was considered an independent biological replicate. Unpaired two-tailed Student’s *t* test. (J) Representative immunofluorescence images of MCT1, GLUT1, and CD31 in the visual cortex of control and *Mctl* ^1ECKO^ mice. Scale bar, 100 μm. (K and L) Quantification of relative endothelial MCT1 (K) and GLUT1 (L) fluorescence intensity in control and *Mct1* ^iECKO^ mice. n = 4 mice per group, with two sections analyzed per mouse. Unpaired two-tailed Student’s *t* test. (M) Representative immunofluorescence images of TGF-β1 and CD31 in the cerebral cortex of control and *Mctl* ^1ECKO^ mice. Scale bar, 100 μm. (N) Quantification of relative endothelial TGF-β1 fluorescence intensity in control and *Mct1* ^iECKO^ mice. n = 3 mice per group, with two sections analyzed per mouse. Unpaired two-tailed Student’s *t* test. (O) Quantification of brain lactate concentrations, normalized to total protein (nmol/mg protein), in control and endothelial-specific *Mct1* conditional knockout (*Mct1* ^iECKO^) mice. n = 7 control and n = 3 *Mct1* ^iECKO^ mice. Unpaired two-tailed Student’s *t* test. Data are presented as mean ± SEM; \**p* < 0.05, \*\**p* < 0.01, and \*\*\**p* < 0.001. See also Figure S8.

We next examined whether endothelial MCT1 deletion, which impaired early cerebrovascular growth, also disrupts the TGF-β1‒MCT1‒lactate signaling axis identified above. Endothelial *Mct1* deletion markedly reduced MCT1 expression and was accompanied by a reciprocal increase in GLUT1 expression (Figures 5J-L), further supporting a role for MCT1 in maintaining the MCT1-high/GLUT1-low endothelial state during early postnatal development. Notably, endothelial TGF-β1 expression was also markedly reduced in endothelial-specific *Mct1* knockout mice (Figures 5M and 5N), consistent with the autoregulatory feedback model proposed in Figure 4. Brain lactate levels were also significantly decreased following endothelial *Mct1* deletion (Figure 5O), further supporting a link between endothelial MCT1 and lactate homeostasis during early postnatal development. Together, these findings indicate that both endothelial MCT1 and canonical TGF-β signaling are required for early postnatal cerebrovascular angiogenesis and support a model in which TGF-β signaling promotes developmental angiogenesis, at least in part, through the endothelial MCT1‒lactate axis. In contrast to the detrimental effects of sustained endothelial MCT1 expression during later cerebrovascular maturation (Figure 2), these findings highlight the stage-dependent regulation of endothelial MCT1 in early vascular growth.

### Astrocytic MCT1 expression attenuates the endothelial TGF-β1‒MCT1 positive feedback loop

In contrast to the marked developmental downregulation of endothelial MCT1 during the first postnatal month, total visual cortex MCT1 protein levels progressively increased over the same period (Figures S2C and S2D). This apparent discrepancy prompted us to ask whether MCT1 expression shifts to another cell type during postnatal development and whether this transition may be associated with the developmental decline in endothelial TGF-β signaling. Because perivascular astrocytes undergo extensive maturation after the first postnatal week^48^, we considered astrocytic MCT1 as a potential regulator of perivascular lactate availability during this developmental transition. Among the monocarboxylate transporters expressed in astrocytes, MCT4 primarily mediates lactate export, whereas MCT1 is expressed in astrocytic perivascular endfeet surrounding brain microvessels and, owing to its higher affinity for lactate, may contribute to lactate uptake^49‒51^. We therefore asked whether astrocyte maturation and astrocytic MCT1 contribute to the developmental attenuation of endothelial TGF-β1‒MCT1 signaling.

We first examined the developmental expression pattern of astrocytic MCT1. Although initial MCT1 immunostaining at P14 appeared less prominent than the endothelial MCT1 signal observed at P7, higher-magnification imaging combined with co-immunostaining for the astrocyte marker S100β revealed previously unrecognized MCT1 signals colocalizing with perivascular astrocyte. Under normal developmental conditions, astrocytic MCT1 expression progressively increased from P7 to P14, coinciding with astrocyte maturation and the decline in endothelial MCT1 expression (Figures 6A and 6B). We next investigated whether this developmental increase in astrocytic MCT1 was influenced by sensory experience (Figure 6C). Dark rearing markedly reduced neuronal activity, as evidenced by decreased c-Fos expression (Figures S2E and S2F), and was accompanied by reduced GFAP expression, consistent with attenuated astrocyte maturation (Figures 6D and 6E). Remarkably, the developmental increase in astrocytic MCT1 observed under normal housing conditions was largely absent in dark-reared mice. Instead, MCT1 expression remained predominantly associated with brain endothelial cells (Figures 6F and 6G), consistent with our previous finding that dark rearing prevents the developmental decline in endothelial MCT1 expression (Figures 1J-L). Together, these findings suggest that sensory experience promotes astrocyte maturation and the developmental increase in astrocytic MCT1 expression, concomitant with the decline in endothelial MCT1.

**Figure 6.**
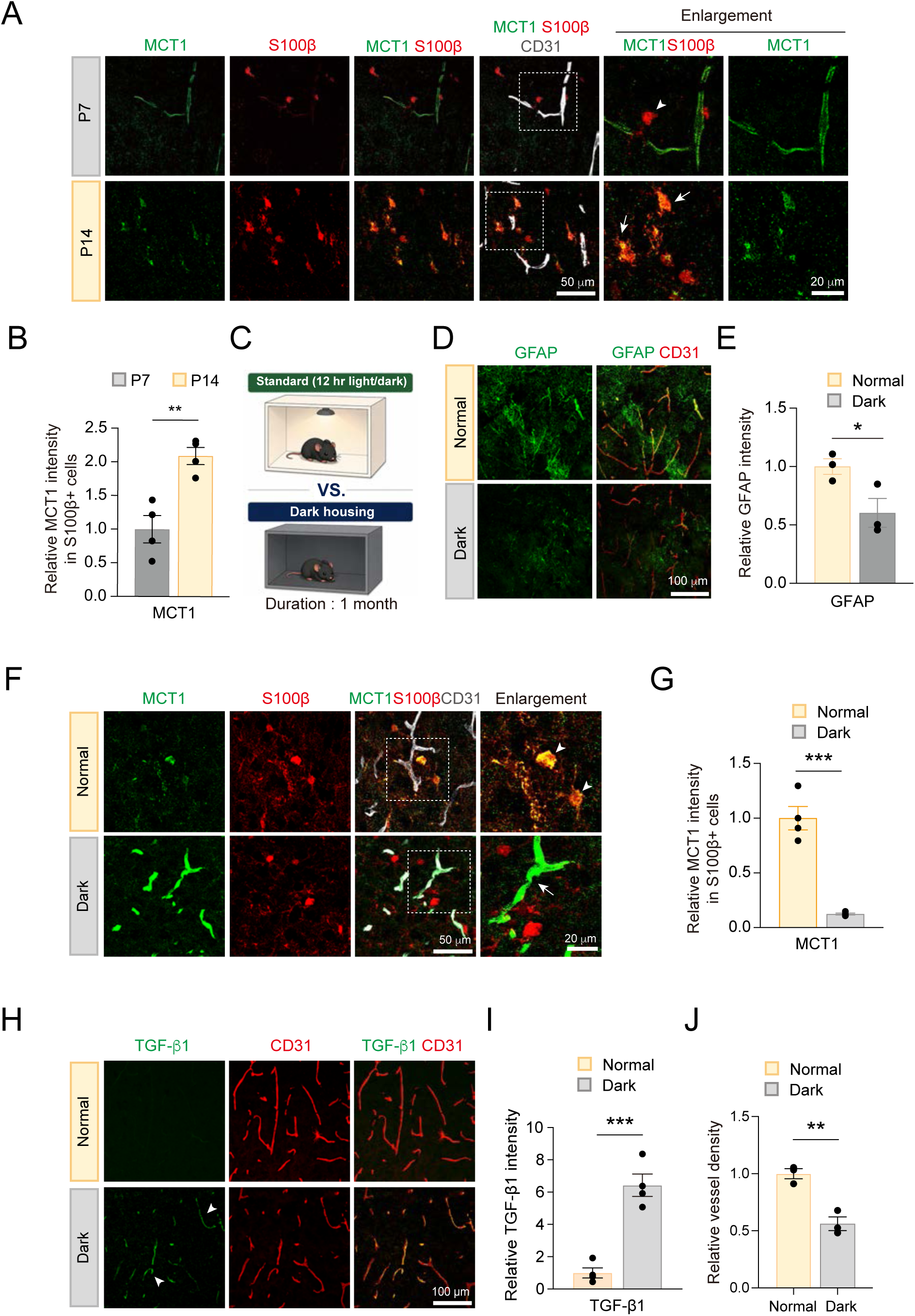
Visual experience promotes the developmental increase in astrocytic MCT1 expression. (A) Representative immunofluorescence images of MCT1, S100β, and CD31 in the visual cortex at P7 and P14. Arrowheads and arrows indicate MCT1-negative and MCT1-positive astrocytes, respectively. Boxed regions are shown at higher magnification. Scale bars, 50 μm and 20 μm (enlargement). (B) Quantification of relative MCT1 fluorescence intensity in S100β-positive astrocytes at P7 and P14. n = 4 mice per age, with two sections analyzed per mouse. Unpaired two-tailed Student’s *t* test (C) Experimental scheme for visual deprivation. Mice were maintained under standard 12-h light/12-h dark conditions (Normal) or in complete darkness (Dark) for 4 weeks. (D) Representative immunofluorescence images of GFAP and CD31 in the visual cortex of normally reared and dark-reared mice. Scale bar, 100 μm. (E) Quantification of relative GFAP expression in normally reared and dark-reared mice. n = 3 mice per group, with two sections analyzed per mouse. Unpaired two-tailed Student’s t test. (F) Representative immunofluorescence images of MCT1, S100β, and CD31 in the visual cortex of normally reared and dark-reared mice. Boxed regions are shown at higher magnification. Arrowheads indicate MCT1-positive, S100β-positive astrocytes, whereas the arrow indicates an MCT1-positive vessel. Scale bars, 50 μm and 20 μm (enlargement). (G) Quantification of relative MCT1 fluorescence intensity in S100β-positive astrocytes from normally reared and dark-reared mice. n = 4 mice per group, with two sections analyzed per mouse. Unpaired two-tailed Student’s *t* test. (H) Representative immunofluorescence images of TGF-β1 and CD31 in the visual cortex of normally reared and dark-reared mice. Arrowheads indicate vascular TGF-β1 immunoreactivity. n = 4 mice per group. Scale bar, 100 μm. (I) Quantification of relative endothelial TGF-β1 fluorescence intensity in normally reared and dark-reared mice. n = 4 mice per group, with two sections analyzed per mouse. Unpaired two-tailed Student’s *t* test. (J) Quantification of vessel density in the visual cortex of normally reared and dark-reared mice. n = 3 mice per group, with ten sections analyzed and averaged per mouse. Each mouse was considered an independent biological replicate. Unpaired two-tailed Student’s *t* test. Data are presented as mean ± SEM; \**p* < 0.05, \*\**p* < 0.01, and \*\*\**p* < 0.001.

We next asked whether this altered MCT1 expression pattern was associated with persistent endothelial TGF-β1 signaling. Endothelial TGF-β1 expression, which normally declined during postnatal development (Figures 3N and 3O), remained elevated in dark-reared mice (Figures 6H and 6I). Dark-reared mice also displayed impaired cerebrovascular maturation, characterized by reduced vascular density and increased BBB permeability (Figure 6J and Figure S3D). Together, these findings suggest that impaired astrocyte maturation and insufficient astrocytic MCT1 upregulation contribute to sustained endothelial TGF-β1‒MCT1 signaling and impaired cerebrovascular maturation under sensory deprivation.

These observations raised the possibility that astrocytic MCT1 contributes to the developmental attenuation of the endothelial TGF-β1‒MCT1 program. To directly test the functional contribution of astrocytic MCT1, we generated tamoxifen-inducible astrocyte-specific *Mct1* conditional knockout mice using the *Aldh1l1-CreERT2* driver (Figure 7A). Because most homozygous knockout mice failed to survive beyond P14, subsequent analyses were performed using heterozygous conditional knockout littermates, which exhibited normal growth during this period. Astrocyte-specific heterozygous *Mct1* deletion (Astro-MCT1 cHet) disrupted the normal developmental transition in endothelial transporter expression, with endothelial MCT1 remaining significantly elevated and GLUT1 reciprocally reduced (Figures 7B-D). The developmental decline in endothelial TGF-β1 normally observed by P14 was also markedly attenuated in astrocytic *Mct1* heterozygous mice (Figures 7E and 7F). Moreover, these mice exhibited impaired cerebrovascular maturation, including reduced vascular density and increased permeability to the low-molecular-weight tracer cadaverine (Figures 7G and 7H). However, no detectable extravascular IgG leakage was observed (Figure 7I).

**Figure 7.**
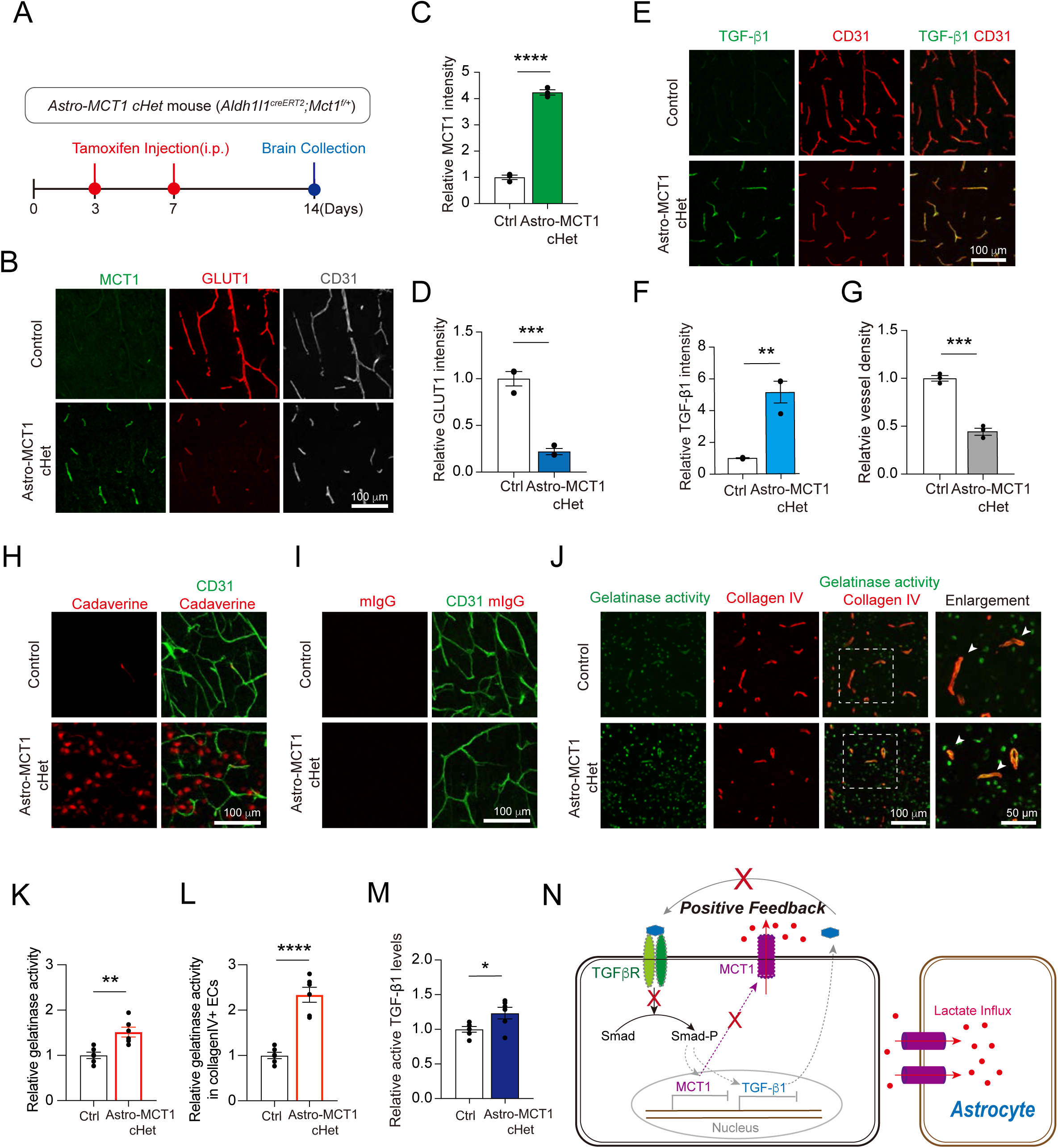
Astrocytic MCT1 attenuates endothelial TGF-β‒MCT1‒lactate signaling to promote vascular maturation. (A) Experimental scheme for inducible astrocyte-specific heterozygous deletion of *Mct1* (*Aldh1l1^creERT^*^2^*;Mct1^f/+^*). Tamoxifen was administered intraperitoneally at P3 and P7, and brains were collected at P14. (B) Representative immunofluorescence images of MCT1, GLUT1, and CD31 in the visual cortex of control and astrocyte-specific conditional *Mctl* heterozygous (Astro-MCT1 cHet) mice. Scale bar, 100 μm. (C and D) Quantification of relative endothelial MCT1 (C) and GLUT1 (D) fluorescence intensity in control and Astro-MCT1 cHet mice. For both MCT1 and GLUT1 analyses, n = 3 mice per group, with two sections analyzed per mouse. Unpaired two-tailed Student’s *t* test. (E) Representative immunofluorescence images of TGF-β1 and CD31 in the visual cortex of control and Astro-MCT1 cHet mice. Scale bar, 100 μm. (F) Quantification of relative endothelial TGF-β1 fluorescence intensity in control and Astro-MCT1 cHet mice. n = 3 mice per group, with two sections analyzed per mouse. Unpaired two-tailed Student’s *t* test. (G) Quantification of vessel density in the visual cortex of control and Astro-MCT1 cHet mice. n = 3 mice per group, with ten sections analyzed and averaged per mouse. Each mouse was considered an independent biological replicate. Unpaired two-tailed Student’s *t* test. (H) Representative images of BBB permeability assessed using cadaverine in control and Astro-MCT1 cHet mice. Similar findings were observed in three mice per group. Scale bar, 100 μm. (I) Representative images of mouse IgG (mIgG) extravasation in the visual cortex of control and Astro-MCT1 cHet mice. Similar findings were observed in three mice per group. Scale bar, 100 μm. (J) Representative images of gelatinase activity and Collagen IV in the cerebral cortex of Astro-MCT1 cHet mice. Boxed regions are shown at higher magnification. Arrowheads indicate gelatinase activity associated with blood vessels. Scale bars, 100 μm (overview) and 50 μm (enlargement). (K) Quantification of relative gelatinase activity fluorescence intensity in control and Astro-MCT1 cHet mice. n = 6 mice per group, with two sections analyzed per mouse. Unpaired two-tailed Student’s *t* test. (L) Quantification of DQ-gelatin fluorescence intensity in collagen IV-positive blood vessels in control and Astro-MCT1 cHet mice. n = 6 mice per group, with two sections analyzed per mouse. Unpaired two-tailed Student’s *t* test. (M) Quantification of relative active TGF-β1 levels in microvessels isolated from the visual cortex of control and Astro-MCT1 cHet mice. n = 6 mice per group. Unpaired two-tailed Student’s *t* test. (N) Proposed model in which astrocytic MCT1-mediated lactate uptake limits extracellular lactate availability and disrupts the endothelial TGF-β-MCT1-lactate positive-feedback loop during cerebrovascular maturation. Data are presented as mean ± SEM; \**p* < 0.05, \*\**p* < 0.01, \*\*\**p* < 0.001, and \*\*\*\**p* < 0.0001.

Because reduced astrocytic MCT1 expression was expected to alter the perivascular microenvironment, potentially increasing extracellular lactate levels, as observed following endothelial MCT1 overexpression (Figures 4M-Q), we examined gelatinase activity and active TGF-β1 levels as potential indicators of changes in extracellular proteolysis and TGF-β1 activation associated with lactate/H^+^ transport.

*In situ* gelatinase assays revealed increased overall gelatinase activity in Astro-MCT1 cHet mice (Figures 7J and 7K), with a particularly marked increase within collagen IV-positive vascular regions (Figures 7J and 7L). Consistent with these changes, ELISA analysis revealed increased levels of active TGF-β1 in brain tissue from Astro-MCT1 cHet mice (Figure 7M). These findings indicate that reduced astrocytic MCT1 expression is accompanied by enhanced perivascular proteolytic activity and increased active TGF-β1 levels, consistent with a role for astrocytic MCT1 in regulating the perivascular environment. Together, these findings indicate that astrocytic MCT1 contributes to the developmental attenuation of endothelial TGF-β1‒MCT1 signaling and the acquisition of mature cerebrovascular properties. Based on these findings, we propose a model in which the maturation-dependent increase in perivascular astrocytic MCT1 may alter local lactate availability, thereby limiting lactate-dependent reinforcement of endothelial TGF-β1 signaling. Attenuation of this positive feedback loop would, in turn, facilitate the developmental downregulation of endothelial MCT1 and promote cerebrovascular maturation (Figure 7N).

## Discussion

During postnatal brain development, cerebral blood vessels transition from an actively growing angiogenic state to a mature state with specialized BBB properties^3‒5,7,52,53^. A central unanswered question is how the timing of this transition is coordinated with the maturation of neuronal circuits and the onset of sensory experience. Here, we show that visual experience drives an endothelial MCT1-to-GLUT1 transition required for cerebrovascular and BBB maturation. Endothelial TGF-β1 signaling sustains the early MCT1-high/GLUT1-low state through an MCT1‒lactate-dependent positive-feedback loop. Around eye opening, sensory activity promotes astrocyte maturation and MCT1 upregulation, potentially limiting perivascular lactate availability and attenuating endothelial TGF-β1 signaling. Together, our findings identify a neuron‒astrocyte‒endothelial signaling mechanism linking sensory experience to endothelial metabolic reprogramming and BBB maturation.

### Neuronal activity coordinates the developmental timing of endothelial MCT1 expression

Although neuronal activity has long been recognized as an important regulator of developmental angiogenesis, as demonstrated in the sensory cortex^4,8^, the molecular mechanisms linking neuronal activity to endothelial maturation remain poorly understood. Recent studies using the retinal vasculature, which shares many structural and functional characteristics with the brain vasculature, have shown that neuronal activity regulates vascular development and barrier formation through pathways such as VEGF and Norrin/β-catenin signaling^2,54^. In addition, transcriptomic analyses of adult brain endothelial cells have identified activity-responsive endothelial gene programs associated with BBB transport regulation^55^. Together, these findings suggest that neuronal activity may influence cerebrovascular maturation through multiple endothelial regulatory mechanisms, potentially including the control of metabolic transporter expression.

Developmental changes in endothelial MCT1 and GLUT1 have been associated with the postnatal shift in cerebral substrate utilization from lactate and ketone bodies toward glucose^16,17,56,57^. However, whether this metabolic transporter transition merely reflects changing developmental demands or actively contributes to cerebrovascular maturation has remained unclear. Our findings indicate that the transition is actively regulated by neuronal activity (Figure 1) rather than being solely a passive adaptation to changing substrate availability. Both visual deprivation and pharmacological suppression of neuronal activity both attenuated the developmental decline in endothelial MCT1 and the reciprocal increase in GLUT1, suggesting that physiological sensory experience helps determine the timing of this metabolic transition. Interestingly, a previous study using a DREADD-based model of glutamatergic neuronal silencing reported an apparently opposite transporter response, with neuronal silencing reducing endothelial MCT1 expression while increasing GLUT1 expression^55^. The basis for this discrepancy remains unclear, but may reflect differences in the mode and duration of neuronal activity manipulation, developmental stage, or brain region examined as previous studies have shown that neuronal activity can exert context-dependent effects on cerebrovascular development^4,6‒8,58‒60^

Beyond its role in substrate transport, MCT1 may contribute to angiogenesis through lactate-dependent signaling. Endothelial metabolism is increasingly recognized as an active regulator of vessel sprouting, with glycolysis supporting endothelial migration and proliferation^61^. Importantly, lactate can function not only as a metabolic substrate but also as a signaling molecule. Previous studies have demonstrated that MCT1-mediated lactate uptake promotes endothelial angiogenic responses through NF-kB/IL-8 and HIF-1 signaling^47,62^. In the brain, lactate signaling through the receptor HCAR1 has also been implicated in VEGF induction and angiogenesis^63^. Beyond signaling, lactate may support the energetic and biosynthetic demands of growing cerebrovascular tissues, particularly during early postnatal development, when circulating lactate and ketone bodies are abundant^17,64,65^.

Importantly, our genetic and viral manipulation studies indicate that the developmental regulation of MCT1 expression, rather than simply its expression level, is critical for cerebrovascular development. Endothelial *Mct1* deletion during early postnatal development impaired vascular growth, consistent with our *in vitro* findings that MCT1-dependent lactate signaling promotes endothelial migration and tube formation (Figure 5 and Figure S8). In contrast, sustained endothelial MCT1 expression beyond its normal developmental window disrupted vascular development and BBB maturation (Figure 2). Together, these findings suggest that endothelial MCT1 supports angiogenic growth during early development but must subsequently be downregulated to permit proper cerebrovascular maturation.

### TGF-β‒MCT1‒lactate signaling regulates the transition from angiogenic growth to endothelial maturation

Our findings identify TGF-β signaling as a key regulator of the developmental MCT1-to-GLUT1 transition (Figures 3 and 4). The underlying mechanism involves an endothelial TGF-β1‒MCT1‒lactate positive-feedback loop that sustains the early MCT1-high state, while its developmental attenuation facilitates the transition to an MCT1-low/GLUT1-high state. This relationship highlights the context-dependent role of TGF-β signaling in endothelial metabolic regulation. Although TGF-β has been reported to promote GLUT1 expression in several non-endothelial and pathological contexts^25,26^, our findings indicate that canonical TGF-β signaling in developing brain endothelial cells instead favors an MCT1-high/GLUT1-low state. Therefore, these findings uncover a previously unrecognized relationship between TGF-β signaling and MCT1-dependent lactate metabolism during cerebrovascular development.

The temporal restriction of this circuit appears to be critical for its developmental function. Deletion of either endothelial *Mct1* or *Tgfbr2* impaired early postnatal vascular growth (Figures 4 and 5), indicating that both MCT1 and TGF-β receptor signaling contribute to developmental angiogenesis. In contrast, sustained MCT1 expression beyond its normal developmental window impaired subsequent vascular and BBB maturation (Figure 2). These findings suggest that TGF-β‒MCT1 signaling supports early vascular growth but must be attenuated to allow subsequent maturation. Although the BBB acquires functional barrier properties during embryonic development, in part through the suppression of endothelial transcytosis ^34^, its tight junction components undergo further molecular maturation during the postnatal period ^66,67^. In this developmental window, sustained endothelial MCT1 expression increased permeability to low-molecular-weight tracers and reduced the tight junction proteins ZO-1 and Claudin-5, whereas no detectable leakage of endogenous IgG or Evans Blue was observed. These findings suggest that persistent MCT1 signaling may preferentially impair maturation of the paracellular barrier. Notably, TGF-β signaling can exert dual, context-dependent effects on BBB integrity, contributing to barrier development and maintenance while also being associated with BBB dysfunction^20,68^. Interestingly, a recent study demonstrated that TGF-β1 promotes BBB leakage by increasing endothelial transcytosis during aging^69^. Unlike this aging-associated transcellular barrier dysfunction, our findings in the developing visual cortex suggest that sustained endothelial MCT1 expression preferentially affects tight junction-associated barrier maturation. Therefore, these observations suggest that TGF-β signaling may regulate BBB function through distinct mechanisms depending on developmental stage and physiological condition, highlighting the importance of timely attenuation of endothelial TGF-β1‒MCT1 signaling during postnatal BBB maturation.

The precise mechanism by which lactate reinforces endothelial TGF-β signaling remains unresolved. One possibility is that lactate regulates TGF-β1 expression through intracellular metabolic signaling. Alternatively, because MCT1 cotransports lactate and protons, changes in MCT1-dependent lactate transport may alter local extracellular pH, thereby potentially facilitating the activation of latent TGF-β1. Previous studies have shown that lactic acid‒induced extracellular acidification can promote TGF-β activation in pulmonary fibrosis^70^. However, whether a similar mechanism operates in developing brain endothelium remains to be determined. Moreover, our observation that sustained endothelial MCT1 expression increased gelatinase activity while reducing collagen IV expression raises the possibility that extracellular matrix remodeling may also contribute to the regulation of TGF-β bioavailability or activation^46,71^.

### Astrocytic MCT1 attenuates endothelial lactate-dependent signaling during postnatal development

Our data suggest that astrocytes provide a mechanism for attenuating the endothelial TGF-β‒MCT1‒lactate circuit (Figure 6 and 7). During early postnatal development, astrocytes undergo extensive morphological and functional maturation, including the formation of perivascular endfeet^48^. As endothelial MCT1 expression declined during postnatal development, MCT1 expression increased in perivascular astrocytes. This developmental increase in astrocytic MCT1 is consistent with previous reports of age-dependent changes in MCT1 expression and cellular distribution in the immature brain^72^. Visual deprivation attenuated the developmental increase in astrocytic MCT1 and was accompanied by persistent endothelial MCT1 and TGF-β1 expression, suggesting that sensory activity coordinates metabolic remodeling across the NVU. Consistent with this model, astrocytic *Mct1* haploinsufficiency sustained endothelial MCT1 and TGF-β1 expression, suppressed GLUT1 induction, and impaired vascular and BBB maturation, alongside increased active TGF-β1 levels and gelatinase activity. These findings provide strong evidence that the developmental increase in astrocytic MCT1 is not merely reciprocal to the decline in endothelial MCT1 but functionally contributes to the attenuation of endothelial TGF-β1-associated signaling and the transition toward vascular maturation.

This mechanism extends the established role of astrocytes in metabolic coupling by implicating astrocytic lactate handling in the regulation of endothelial developmental state. Previous studies have established extensive metabolic interactions between astrocytes and neurons^73^, whereas our findings suggest that astrocytes may also regulate the vascular microenvironment during postnatal maturation. The reciprocal changes in endothelial and astrocytic MCT1 further raise the possibility that MCT1-dependent lactate handling is developmentally redistributed across NVU cell types as the vasculature transitions from growth to maturation. Although other MCT-expressing NVU cell types, including neurons and pericytes, may contribute to local lactate homeostasis^74,75^, our findings suggest that astrocytic MCT1 plays a particularly important role in coordinating perivascular lactate handling with endothelial maturation. However, the mechanisms by which visual experience-driven neuronal activity upregulates astrocytic MCT1 remain unclear. Given that astrocytic MCT1 expression increases markedly around eye opening (∼P14, Figure 6), further characterization of astrocytic gene expression profiles at this developmental stage will be important for identifying the molecular mechanisms underlying this activity-dependent upregulation.

Taken together, our studies support a model in which neuronal activity coordinates cerebrovascular maturation by regulating the timing of an endothelial lactate-dependent angiogenic program and its subsequent transition toward vascular and BBB maturation (Figure 8). More broadly, this work establishes a conceptual link between neuronal activity, intercellular metabolic signaling, and cerebrovascular development, providing a framework for understanding how experience-dependent neurovascular interactions shape brain maturation and function.

**Figure 8.**
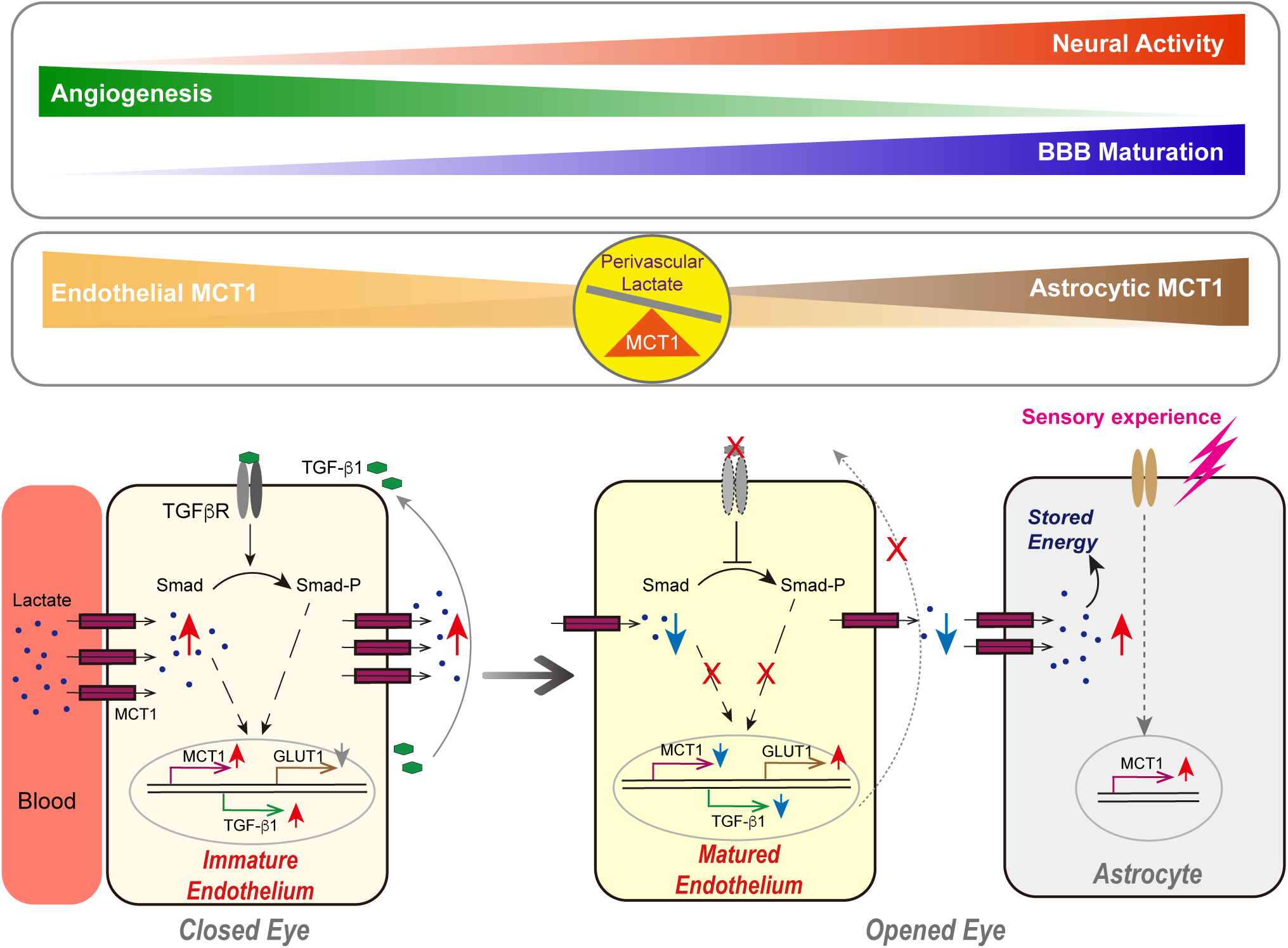
Proposed model of visual experience‒dependent regulation of the transition from angiogenic growth to cerebrovascular and BBB maturation.

## Data availability

The RNA-sequencing data generated in this study are publicly available in the Gene Expression Omnibus (GEO) under accession number GSE178918. All other data supporting the findings of this study and research materials, including mouse models, are available from the corresponding authors upon reasonable request.

## Acknowledgements

We thank Dr. Bin Zhou for providing *Mfsd2a-CreERT2* mice; Dr. Hyungju Park for providing *Aldh1l1-CreERT2* mice; and the Brain Research Core Facilities at the Korea Brain Research Institute (KBRI) for image analysis. This work was supported by the KBRI basic research program of the Korea Brain Research Institute funded by the Ministry of Science and ICT (KBRI 26-BR-01-02 and KBRI 26-BR-05-06), and the Bio & Medical Technology Development Program of the NRF, funded by the Korean government (MSIT) (NRF-2022M3E5E8017701 and RS-2026-25475900) to W.O.; the Young Researcher Program of the NRF, funded by the Korean government (MSIT) (2020R1C1C1010509) to N.K.; and the Global-Learning & Academic Research Institution for Master’s PhD Students and Postdocs (G-LAMP) Program of the NRF, funded by the Ministry of Education (RS-2023-00301850) to S.C.

## Author contributions

WO conceived and designed the project. NK, SC, EC, JJ and HK conducted experiments and acquired data. NK, SC and WO wrote the manuscript.

## Materials and Methods

### Animals and treatments

*Pdgfrb-CreERT2* (JAX stock no. 030201), *Camk2a-CreERT2* (JAX stock no. 012362), *Tie2-Cre* (JAX stock no. 008863), *Tgfb1-flox* mice (JAX stock no. 065809), and RiboTag (Rpl22^HA/HA^; JAX stock no. 011029) mice were purchased from The Jackson Laboratory (Bar Harbor, ME, USA). *Aldh1l1-CreERT2* mice (JAX stock no. 031008) were kindly provided by Dr. Hyungju Park (Korea Brain Research Institute, Daegu, Republic of Korea). *Mfsd2a-CreERT2* mice were kindly provided by Dr. Bin Zhou^76^. *Mct1-flox* mice (Cyagen, S-CKO-05084) were purchased from Cyagen Biosciences (Suzhou, China). *Tgfbr2-flox* mice (KMB resource no. KMB-PX00004915) were purchased from the Laboratory Animal Resource Center, Korea Research Institute of Bioscience and Biotechnology (KRIBB, Cheongju, Republic of Korea). All mice were maintained on a C57BL/6J background (JAX stock no. 000664) and housed under standard conditions with a 12-h light/dark cycle.

For inducible gene deletion, tamoxifen (Sigma-Aldrich, T5648) was dissolved in corn oil (Sigma-Aldrich, C8267) and administered by intraperitoneal (i.p.) injections of 100 mg/kg to mice before experiment. For experiments in which brains were collected at P7, tamoxifen was administered at P3, whereas for experiments in which brains were collected at P14, tamoxifen was administered at P3 and P7. To suppress neuronal activity during the peri-eye-opening period, muscimol (Sigma-Aldrich, G019) was administered by i.p. injection at 0.5 mg/kg once daily from P10 to P14, followed by brain collection at P15. Control mice received the corresponding vehicle using the same administration schedule. For dark-rearing experiments, mice were housed in complete darkness for 4 weeks. Age-matched control mice were maintained under standard light/dark conditions. Animals were analyzed immediately after the dark-rearing period.

All animal experiments were approved by the Institutional Animal Care and Use Committee of the Korea Brain Research Institute (IACUC-24-00046 and IACUC-25-00013) and were performed in accordance with the National Institutes of Health Guide for the Care and Use of Laboratory Animals and ARRIVE guidelines.

### Adeno-associated virus administration

For sustained expression of MCT1 in brain endothelial cells, we used AAV-BI30, an engineered AAV capsid that efficiently transduces endothelial cells throughout the central nervous system^31^. AAV-BI30 vectors expressing either GFP (AAV-GFP) or MCT1-V5 (AAV-MCT1-V5) were administered to neonatal mice via the superficial temporal vein at postnatal day 3 (P3) at a dose of 5 × 10^10^ vector genomes (vg) per animal in a total volume of 1 μL. AAV-GFP was used as the control vector. Brain tissues were collected at the indicated time points for subsequent analyses.

### Cell culture and treatments

The human cerebral microvascular endothelial cell (hCMEC/D3) was purchased from Millipore (SCC066). The hCMEC/D3 cells were cultured in endothelial cell medium (ScienCell, 1001) supplemented with 5% fetal bovine serum (ScienCell, 0025), endothelial cell growth supplement (ScienCell, 1052), 100 U/mL penicillin, and 100 μg/mL streptomycin (ScienCell, 0503) at 37 °C in a humidified atmosphere containing 5% CO_2_. Cell culture dishes (100 mm; Greiner Bio-One, 664160`—`013) were precoated with rat tail collagen type I (5 μg/mL; Millipore, 08-115) at 37 `°`C for at least 1 h. The culture medium was replaced every 2`—`3 days until cells reached 70-80% confluence. Cells were then detached using 0.25% trypsin-EDTA (HyClone, SH30042.02) at 37 °C for 1 min. Trypsinization was terminated by adding serum-containing medium, followed by centrifugation at 196 × g for 3 min. Cells were seeded at a density of 3 × 10^5^ cells per 60-mm dish and allowed to adhere overnight before treatment.

To activate TGF-β signaling, cells were treated with recombinant human TGF-β1 (10 ng/mL; BioLegend, 781802) for the indicated durations (24-72 h). To examine the involvement of TGF-β signaling, cells were treated with the SMAD3 inhibitor SIS3 (1 or 5 μM; Sigma-Aldrich, 566405) or the TGF-β receptor type I kinase inhibitor galunisertib (1 or 5 μM; Sigma-Aldrich, SML2851) for 24 h. For lactate treatment experiments, cells were exposed to sodium L-lactate (10 mM; Sigma-Aldrich, L7022) for 48 h. Control cells received equivalent volumes of the corresponding vehicles (PBS, 0.1% DMSO, or PBS containing 0.1% BSA). To screen signaling factors potentially regulating MCT1 and GLUT1 expression, cells were treated with recombinant human TGF-β1 (10 ng/mL; BioLegend, 781802), IGF-I (100 ng/mL; Sino Biological, 10598-HNAE), IGF-II (100 ng/mL; Sino Biological, 13032-HNAY), VEGF (100 ng/mL; Sino Biological, 11066-HNAH), BDNF (100 ng/mL; Sino Biological, 50240-MNAS), or lithium chloride (LiCl; 20 mM; Sigma-Aldrich, 213233) for 72 h. Control cells received equivalent volumes of the corresponding vehicles (PBS or PBS containing 0.1% BSA).

### Protein sample preparation

Cells were washed with ice-cold phosphate-buffered saline (PBS, pH 7.4) and harvested using a cell scraper (SPL, 90020) in ice-cold mild lysis buffer (100 mM Tris-HCl [pH 7.4], 10 mM EDTA, 150 mM NaCl, and 1% Triton X-100) supplemented with a protease inhibitor cocktail (Thermo Fisher Scientific, 78444). The cell lysates were homogenized using an insulin syringe (BD, 328822), incubated on ice for 20 min, and centrifuged at 12,000 × g for 10 min at 4 °C. For brain tissue protein extraction, mice were perfused with ice-cold PBS, and brain tissues were homogenized on ice in lysis buffer supplemented with protease and phosphatase inhibitors using Taco™ Pre-loaded Steel Bead Tubes (GeneReach, a-psbt) and a Taco™ Prep Bead Beater (GeneReach Biotechnology Corp., Taiwan). The homogenates were incubated on ice for 20 min and subsequently centrifuged at 12,000 × g for 10 min at 4 °C. For cerebral microvessel isolation, mice were perfused with ice-cold PBS, and the visual cortex was dissected. The tissue was homogenized using a glass Dounce homogenizer (Thermo Fisher Scientific, 3432T40). The homogenate was passed through a 40-μm cell strainer (SPL, 93040), and the fraction retained on the strainer was collected for microvessel isolation. The isolated microvessel fraction was lysed in the buffer described above, incubated on ice for 20 min, and centrifuged at 12,000 × g for 10 min at 4 °C. The resulting supernatants were collected, and protein concentrations were determined using the Pierce™ BCA Protein Assay Kit (Thermo Fisher Scientific, 23228).

### Western blotting

For western blot analysis of cell lysates, 50-60 μg of protein was separated by 10-14% SDS-PAGE and transferred onto a polyvinylidene fluoride (PVDF) membrane. For mouse samples, 50 μg of protein from brain homogenates or 5-10 μg of protein from isolated cerebral microvessel fractions was subjected to 10-14% SDS-PAGE and transferred onto PVDF membranes. Protein samples were mixed with 4× sample buffer [250 mM Tris-HCl (pH 6.8), 8% sodium dodecyl sulfate, 0.1% bromophenol blue, 40% glycerol, and 100 mM DTT]. Membranes were blocked with 5% skim milk or 5% bovine serum albumin (BSA) in Tris-buffered saline containing 0.1% Tween-20 (TBS-T) for 1 h and incubated overnight at 4 °C with the indicated primary antibodies diluted in the corresponding blocking buffer. The primary antibodies used were anti-TGF-β1 (1:1,000, Cell Signaling Technology, 3711S), anti-p-SMAD2 (1:1,000, Cell Signaling Technology, 18338S), anti-SMAD2/3 (1:1,000, Santa Cruz Biotechnology, sc-133098), anti-HIF-1α (D1S7W) (1:1,000, Cell Signaling Technology, 36169), anti-MCT1 (1:1,000, Cell Signaling Technology, 85680), anti-GLUT1 (1:1,000, Cell Signaling Technology, 12939S), anti-CD31 (1:1,000, R&D Systems, AF3628), anti-ZO-1 (1:1,000, Thermo Fisher Scientific, 61-7300), anti-occludin (1:1,000, Thermo Fisher Scientific, 71-1500), anti-claudin-5 (1:1,000, Thermo Fisher Scientific, 34-1600), anti-caveolin-1 (1:1,000, Abcam, ab2910), anti-V5-tag (1:1,000, Cell Signaling Technology, 13202), anti-GFP (1:1,000, Santa Cruz Biotechnology, sc-9996), and anti-β-actin (1:5,000, Cell Signaling Technology, 5125). Following incubation with appropriate horseradish peroxidase (HRP)-conjugated secondary antibodies, protein bands were detected using a Fusion FX chemiluminescence imaging system (Vilber Lourmat, Marne-la-Vallée, France) and quantified using ImageJ software.

### Immunostaining

For immunofluorescence analysis, mouse brains were fixed overnight in 2% paraformaldehyde (PFA) and cryoprotected in 20% sucrose in PBS. Brain tissues were sectioned at a thickness of 20 μm using a cryostat (Leica Microsystems, Germany). Sections were permeabilized with PBS containing 0.2% Triton X-100 for 10 min and blocked in PBS containing 1% bovine serum albumin (BSA), 0.2% Triton X-100, and 0.3 M glycine for 1 h at room temperature. Sections were then incubated overnight at 4 °C with primary antibodies diluted in PBS containing 1% BSA and 0.2% Triton X-100.

The following primary antibodies were used: anti-MCT1 (1:250, Santa Cruz Biotechnology, sc-365501), anti-MCT1 (1:250, Cell Signaling Technology, 85680S), anti-GLUT1 (1:250, Merck Millipore, 07-1401), anti-CD31 (1:150, BD Pharmingen, 553370), anti-GFP (1:500, Abcam, ab13970), anti-DsRed (1:500, Santa Cruz Biotechnology, sc-390909), anti-V5-tag (1:500, Cell Signaling Technology, 13202), anti-V5-tag (1:500, Thermo Fisher Scientific, R960-25), anti-ZO-1 (1:250, Thermo Fisher Scientific, 61-7300), anti-Claudin-5 (1:250, Thermo Fisher Scientific, 35-2500), anti-c-Fos (1:250, Abcam, ab190289), anti-GFAP (1:500, Cell Signaling Technology, 12389), anti-S100 (1:500, Agilent, Z0311), anti-MAL (1:250, Santa Cruz Biotechnology, sc-390687), anti-HSPB1 (1:250, Santa Cruz Biotechnology, sc-13132), and anti-Collagen IV (1:500, Abcam, ab6586), anti-TGF-β1 (1:250, Santa Cruz Biotechnology, sc-130348), anti-TGF-β receptor 2 (TGFBR2; 1:250, Santa Cruz Biotechnology, sc-17792), and anti-phospho-SMAD2 (1:250, Cell Signaling Technology, 18338).

Following primary antibody incubation, sections were washed three times with PBS containing 0.2% Tween-20 and incubated for 1 h at room temperature with Alexa Fluor 488-, 568-, or 647-conjugated secondary antibodies (1:500, Thermo Fisher Scientific). For negative controls, sections were processed under identical conditions without primary antibodies and incubated with secondary antibodies alone. Fluorescence images were acquired using a Nikon Eclipse Ti-U microscope (Nikon, Japan) or a Leica TCS SP8 confocal microscope (Leica Microsystems, Germany). Images were processed and analyzed using ImageJ and Adobe Photoshop CC 2021.

For immunostaining of brain sections used in BBB permeability assays, brains were fixed overnight in 2% PFA and embedded in 4% agarose in PBS. Brain tissues were sectioned at a thickness of 100 μm using a vibratome (VT1000 S, Leica Microsystems, Germany). Free-floating sections were permeabilized with PBS containing 0.2% Triton X-100 for 10 min and blocked in PBS containing 1% BSA, 0.2% Triton X-100, and 0.3 M glycine for 5 h at room temperature. Sections were subsequently incubated overnight at 4 °C with primary antibodies diluted in PBS containing 1% BSA and 0.2% Triton X-100. After primary antibody incubation, sections were washed three times for 30 min each with PBS containing 0.2% Tween-20 and incubated overnight at 4 °C with Alexa Fluor 488-, 568-, or 647-conjugated secondary antibodies (1:500, Thermo Fisher Scientific). Images were acquired using a Leica TCS SP8 confocal microscope (Leica Microsystems, Germany).

### *In situ* zymography

Gelatinase activity in brain sections was assessed by *in situ* zymography using the EnzChek™ Gelatinase/Collagenase Assay Kit (Thermo Fisher Scientific, E12055) according to the manufacturer’s instructions. Briefly, mouse brains were rapidly dissected, immediately frozen in liquid nitrogen, and stored at -80 °C until sectioning. Fresh-frozen brains were sectioned at a thickness of 10 μm using a cryostat. The sections were incubated with 20 μg/mL fluorescein-conjugated DQ gelatin in reaction buffer for 3 h at 37 °C in a humidified chamber. After incubation, the sections were washed with PBS and fixed with 4% paraformaldehyde (PFA) for 10 min. Subsequent immunofluorescence staining was performed as described in the Immunostaining section above. Fluorescence images were acquired using a Nikon Eclipse Ti-U microscope (Nikon, Japan).

### BBB permeability assay

BBB permeability was assessed using Alexa Fluor 555 Cadaverine (Thermo Fisher Scientific, A30677), 3-kDa Texas Red Dextran, Lysine Fixable (Thermo Fisher Scientific, D3328), Evans blue, or endogenous IgG. For fluorescent tracer assays, Alexa Fluor 555 Cadaverine (5 mg/mL) or Texas Red Dextran (15 mg/mL) was diluted fivefold in PBS, and 100 μL of the diluted tracer solution was administered via retro-orbital injection. After 2 h of circulation, mice were transcardially perfused with PBS to remove intravascular tracer, and brains were collected and fixed overnight in 2% paraformaldehyde (PFA). For Evans blue assays, mice were retro-orbitally injected with 100 μL of filtered 2% Evans blue and allowed to circulate for 30 min before PBS perfusion and brain collection. Endogenous IgG leakage was assessed by direct detection of mouse IgG using an Alexa Fluor-conjugated anti-mouse IgG secondary antibody. Tissue processing and immunostaining were performed as described in the Immunostaining section above.

### Transendothelial electrical resistance (TEER) measurements

Cells were seeded at a density of 8,000 cells per insert (Day 0) onto 0.4-μm-pore polyethylene terephthalate (PET) Transwell inserts (BD Falcon, 353095) in 24-well plates. The inserts were precoated with rat tail collagen type I (5 μg/mL) for at least 1 h. The culture medium was replaced every 2-3 days. TEER was measured daily from Day 1 through Day 10 using an EVOM2 epithelial voltohmmeter equipped with STX2 chopstick electrodes (World Precision Instruments, USA). On Day 7 after seeding, after confirming the formation of a confluent monolayer, the culture medium was replaced with fresh medium containing vehicle (control) or recombinant human TGF-β1 (10 ng/mL; BioLegend, 781802). Before measurement, the electrodes were disinfected with 70% ethanol and equilibrated in prewarmed PBS for 5 min. TEER measurements were performed in triplicate using three Transwell inserts per condition, and the mean value was calculated. TEER values were expressed in Ω·cm^2^ after normalization to the membrane surface area (0.33 cm^2^).

### RiboTag-based translatome sequencing and bioinformatic analysis

Ribosome-associated mRNA was isolated from visual cortex microvessels of Tie2-Cre;Rpl22^HA/HA^ mice at P10 and P30 (n = 4 mice per group) by anti-HA immunoprecipitation as described previously^27^. Sequencing libraries were prepared using the NEBNext Single Cell/Low Input RNA Library Prep Kit for Illumina (New England Biolabs, USA) according to the manufacturer’s instructions and sequenced on an Illumina NovaSeq 6000 platform (Macrogen, Korea). Raw reads were aligned to the Mus musculus reference genome (GRCm39) using STAR software (version 2.7.10) with default parameters^77^. After alignment, the number of reads mapped to the gene features (GTF file of GRCm39.112) was counted using HTSeq software^78^. Read counts for the samples in each condition were then normalized using TMM (trimmed mean of M-values) normalization of the edgeR package^79^. The numbers of reads of gene features were converted to log2-values after adding one (pseudo count) to the read counts. To identify Differentially expressed ribosome-associated transcripts, a previously reported statistical hypothesis test was performed^80^. Briefly, for each gene, the t-statistic value was calculated using Student’s *t* test to compare P30 and P10. The empirical distributions of the t-statistic values for the null hypothesis (i.e., no differential expression of the gene) were then estimated by performing all possible random permutations of the samples. The adjusted p-values of each gene for the individual tests were computed using a two-tailed test with the empirical null distribution and combined using Stouffer’s method. The adjusted p-values were then computed with the Storey method. Of the annotated features, 16,955 genes with a unique gene symbol were retained for downstream analysis. Differential changes in the endothelial translatome were identified by adjusted p-values ≤ 0.05 and absolute log2-fold changes > 1. Functional over-representation analysis was performed using ConsensusPathDB^28^. Gene Ontology (GO) terms with *p*-values < 0.05 were considered significantly enriched. RNA sequencing data have been deposited in the Gene Expression Omnibus (GEO) under accession number GSE178918.

### Quantitative reverse transcription PCR (qRT-PCR)

Ribosome-associated endothelial RNA was isolated from the visual cortices of Tie2-Cre;Rpl22^HA/HA^ mice at P10 and P30 by anti-HA immunoprecipitation using the RiboTag approach, as described above^27^. cDNA was synthesized and amplified from 1 ng of RiboTag-isolated endothelial RNA using the NEBNext Single Cell/Low Input cDNA Synthesis & Amplification Module (New England Biolabs, E6421L) according to the manufacturer’s instructions. Quantitative real-time PCR (RT-qPCR) was performed in triplicate using SYBR Green I Master Mix (Roche, 04707516001) on a LightCycler 480 system (Roche). Relative gene expression was calculated using the 2^-ΔΔCt^ method with *Gapdh* as the reference gene. Primer sequences used for RT-PCR and RT-qPCR analyses are listed in Table S2.

### Tube formation assay

Growth Factor Reduced (GFR) Basement Membrane Matrix (Corning, 354230) was thawed overnight at 4 °C and maintained on ice until use. A total of 75 μL of the matrix was added to each well of a prechilled 96-well culture plate using precooled pipette tips. The plate was incubated at room temperature for 60 min to allow the matrix to polymerize. hCMEC/D3 cells (3 `×` 10^4^ cells in 100 μL of culture medium) were seeded into each matrix-coated well and treated with vehicle (0.1% DMSO), AZD3965 (1 μM; MedChemExpress, HY-12750), sodium L-lactate (10 mM; Sigma-Aldrich, L7022), or a combination of AZD3965 and sodium L-lactate, as indicated. For cotreatment experiments, AZD3965 was added 1 h before sodium L-lactate treatment. The plate was incubated for 8 h at 37 °C in a humidified atmosphere containing 5% CO_2_. Tube-like structures were imaged at 20 `×` magnification using a phase-contrast microscope (Nikon Eclipse TS2, Japan). The number of junctions, total tube length, and number of tubes were quantified using the Angiogenesis Analyzer plugin in ImageJ software. Each condition was assessed in triplicate wells.

### Wound healing assay

Cell migration was evaluated using a wound-healing assay. hCMEC/D3 cells (2 × 10^5^ cells per well) were seeded into 12-well plates and cultured until a confluent monolayer was formed. After overnight serum starvation, a linear scratch was made using a sterile 1-mL pipette tip. The cells were washed with PBS to remove detached cells and then cultured in medium containing 0.3% FBS supplemented with vehicle (0.1% DMSO), AZD3965 (1 μM; MedChemExpress, HY-12750), or sodium L-lactate (10 mM; Sigma-Aldrich, L7022). Images of the wound area were acquired at 0 and 24 h using a phase-contrast microscope (Nikon Eclipse TS2, Tokyo, Japan). Wound closure was quantified as the percentage reduction in wound area at 24 h relative to the initial wound area at 0 h using ImageJ software. Experiments were performed in triplicate for each condition.

### Colorimetric lactate assay

L-Lactate concentrations in cell and tissue lysates were measured using a colorimetric lactate assay kit (Cell Biolabs, Inc., MET-5012) according to the manufacturer’s instructions. For cell lysate preparation, cells were washed with ice-cold phosphate-buffered saline (PBS, pH 7.4), harvested in 100 μL of 1× lactate assay buffer using a cell scraper (SPL, 90020), and homogenized using an insulin syringe (BD, 328822). The lysates were centrifuged at 10,000 × g for 5 min at 4 °C. For tissue lysate preparation, mice were perfused with ice-cold PBS, and fresh brain tissues were immediately collected and homogenized in 1× lactate assay buffer using Taco™ Pre-loaded Steel Bead Tubes (GeneReach, a-psbt) and a Taco™ Prep Bead Beater. The homogenates were centrifuged at 12,000 × g for 10 min at 4 °C. The resulting supernatants were transferred to Amicon® Ultra 0.5-mL centrifugal filters (10-kDa molecular weight cutoff [MWCO]; Millipore, UFC501096) to remove lactate dehydrogenase (LDH) and centrifuged again at 12,000 × g for 10 min. Samples and lactate standards were incubated at 37 °C for 30 min, and absorbance was measured at 570 nm. Lactate concentrations were calculated from a standard curve and normalized to total protein content.

### Enzyme-linked immunosorbent assay (ELISA) for free active TGF-β1

To measure free active TGF-β1 levels, mice were perfused with ice-cold PBS, and microvessels were isolated from the dissected visual cortex. The isolated microvessels were homogenized in lysis buffer (20 mM Tris-HCl [pH 7.4], 150 mM NaCl, 2 mM EDTA, and 0.5% Triton X-100) supplemented with a protease inhibitor cocktail. The lysates were incubated on ice for 10 min and centrifuged at 12,000 × g for 10 min at 4 °C. The resulting supernatants were collected and stored at -80 °C until analysis. Total protein concentrations were determined using the Pierce™ BCA Protein Assay Kit (Thermo Fisher Scientific, 23228). Free active TGF-β1 levels were measured using the LEGEND MAX™ Free Active TGF-β1 ELISA Kit (BioLegend, 437707) according to the manufacturer’s instructions. Absorbance was measured at 450 nm with wavelength correction at 570 nm. Free active TGF-β1 concentrations were calculated from a standard curve and normalized to the total protein content of each sample.

### Statistical analysis

All values are presented as the mean ± standard error of the mean (SEM). For histological analyses, measurements from multiple sections obtained from the same mouse were averaged to generate a single value per animal, and each mouse was considered an independent biological replicate for statistical analysis. Specifically, for quantification of vessel density, vessel diameter, and vessel length, ten sections per mouse were analyzed, and the measurements were averaged for each mouse before statistical analysis. A two-tailed Student’s *t* test or Welch’s *t* test was used to compare two independent groups, as appropriate. For comparisons involving multiple groups or factors, one-way or two-way analysis of variance (ANOVA) was performed, as appropriate, followed by Tukey’s, Dunnett’s, or Sidak’s multiple-comparisons test. All statistical analyses were performed using GraphPad Prism 11.0.0 (GraphPad Software, Inc.). p < 0.05 was considered statistically significant.

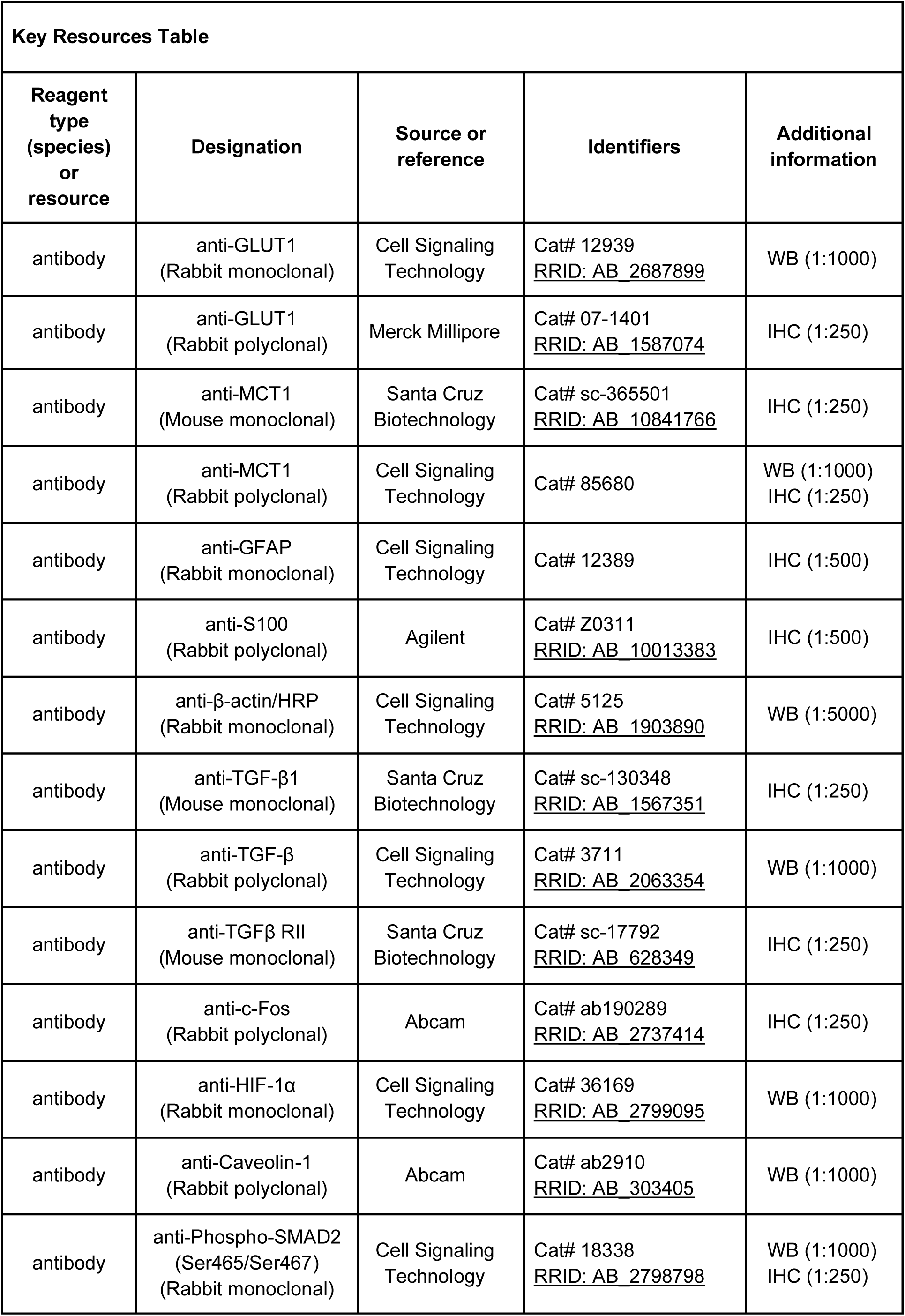
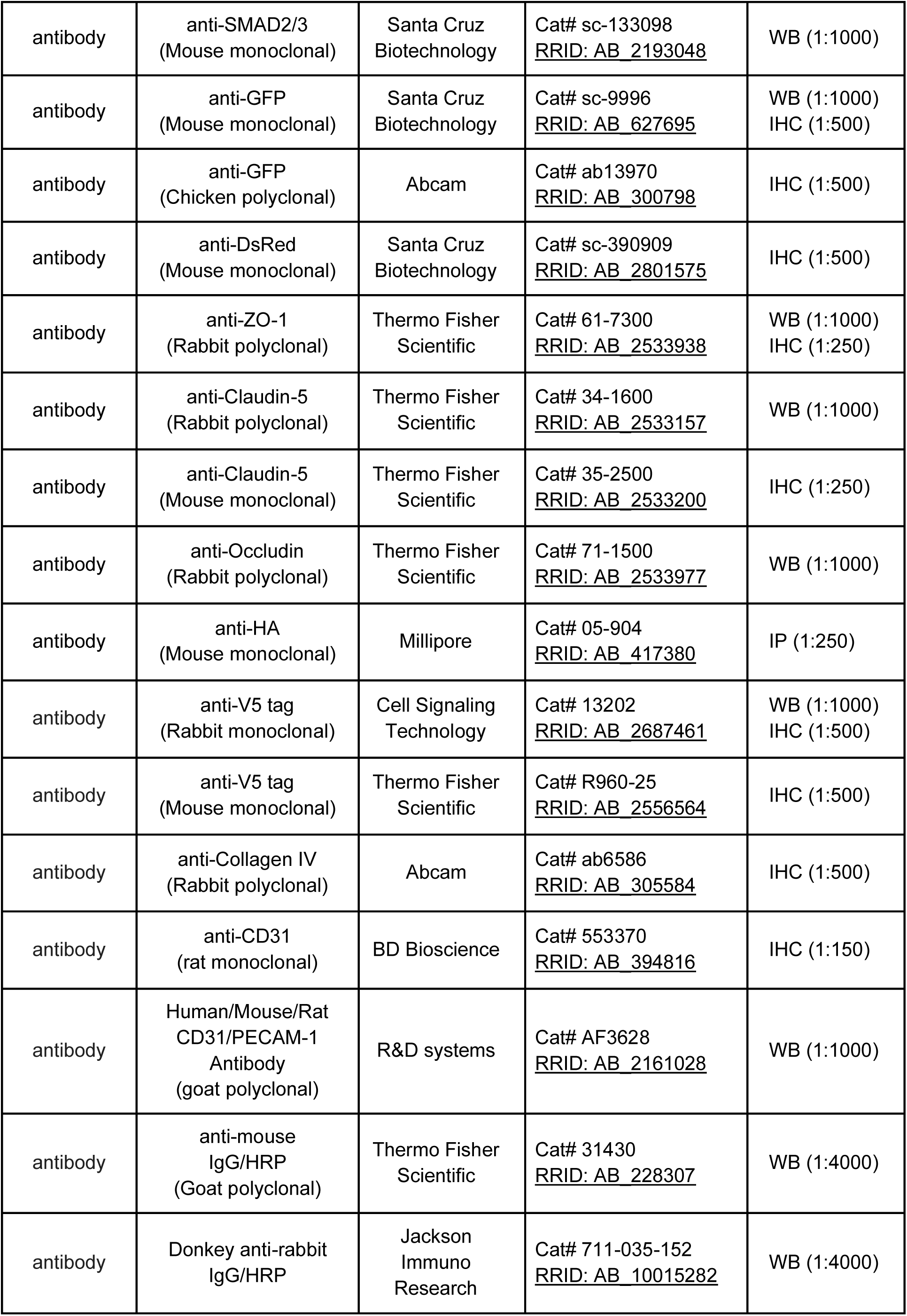
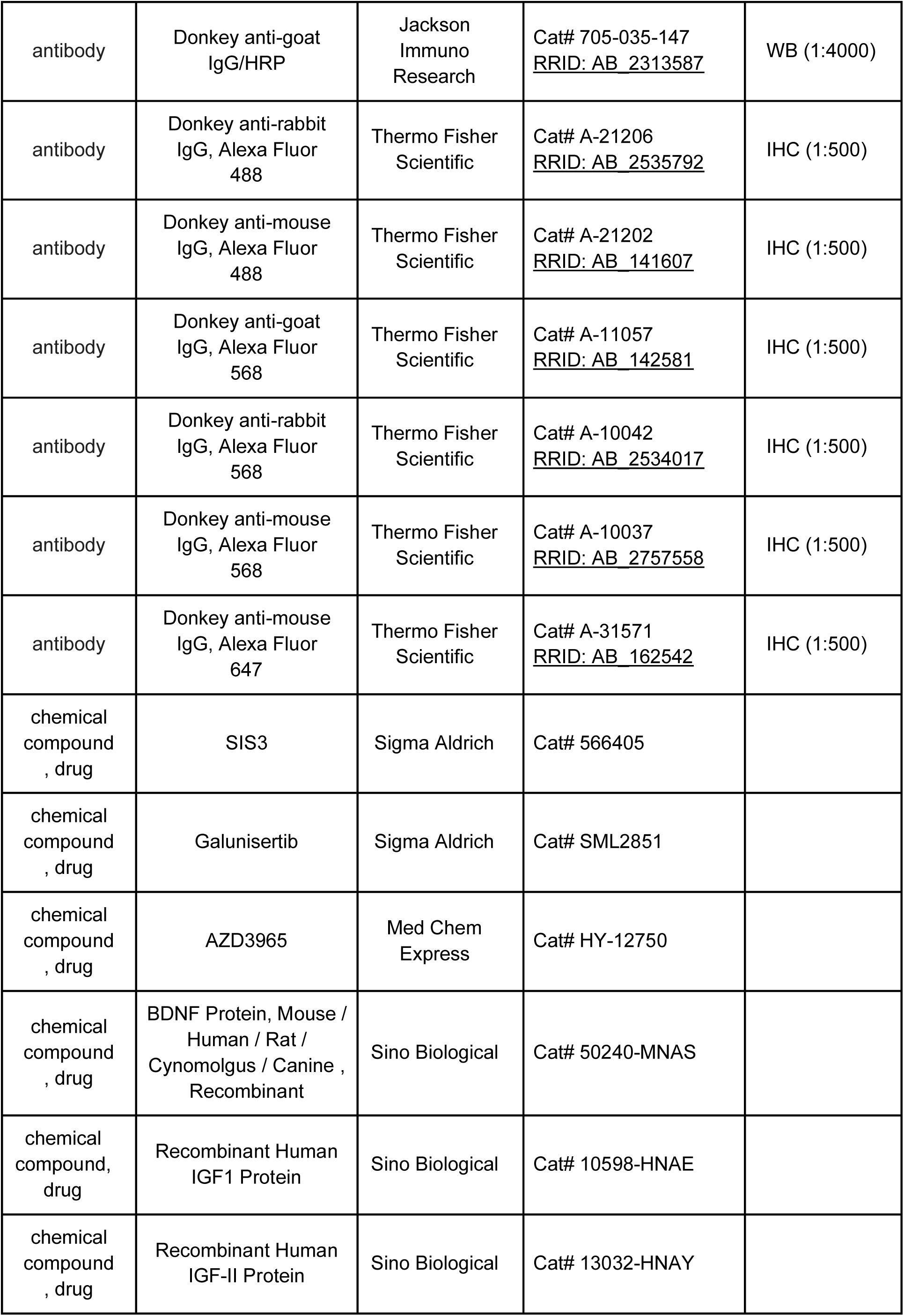
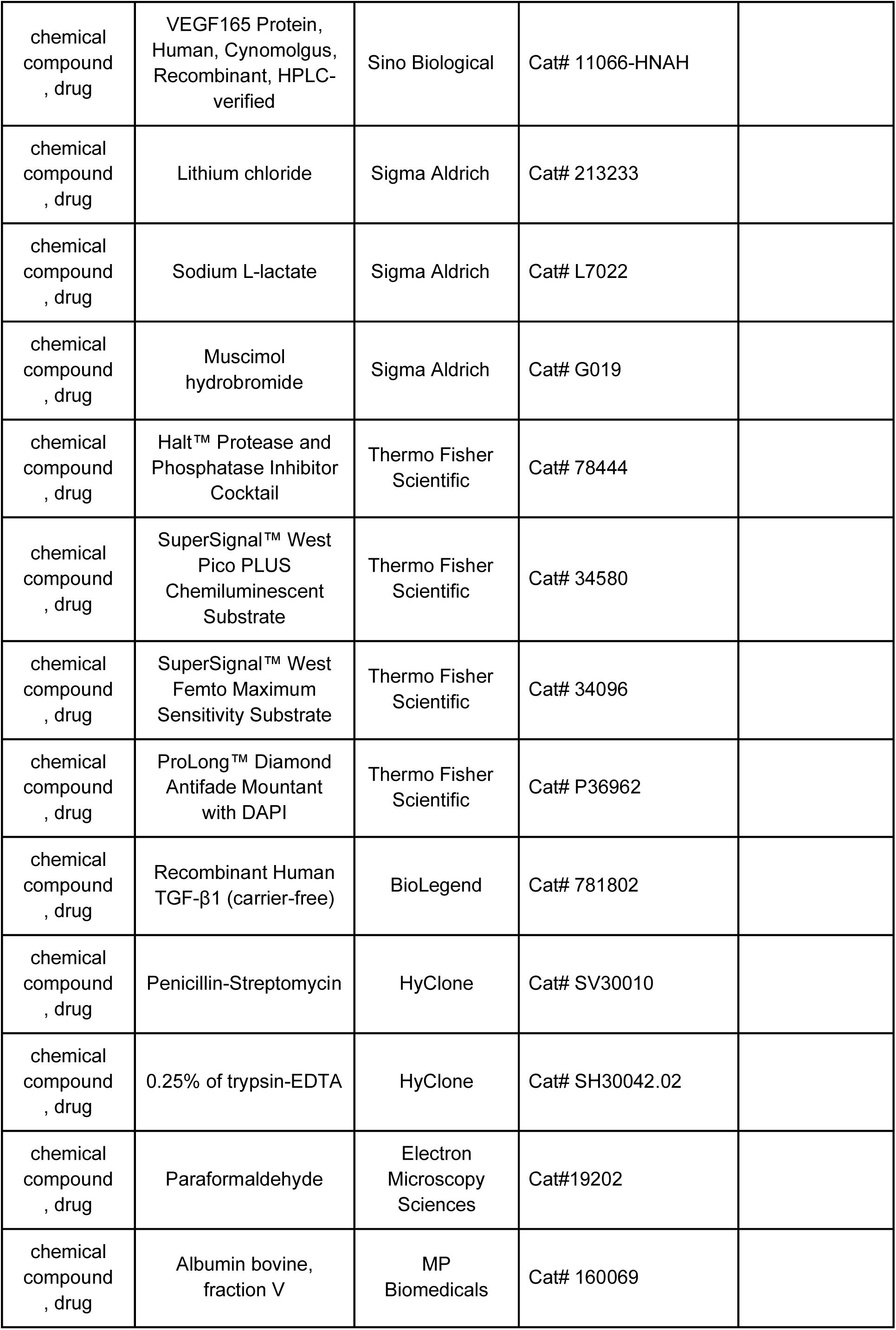
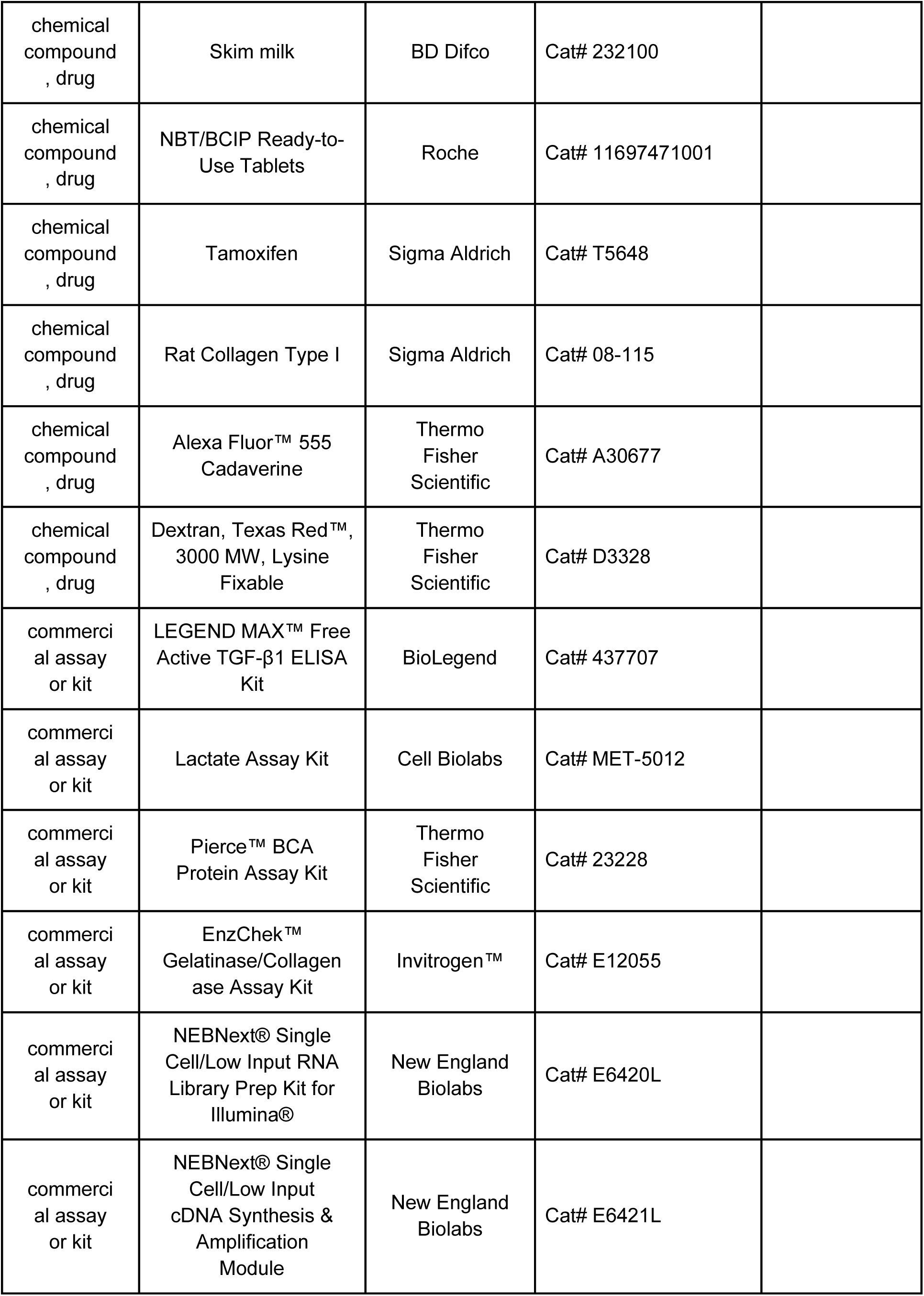
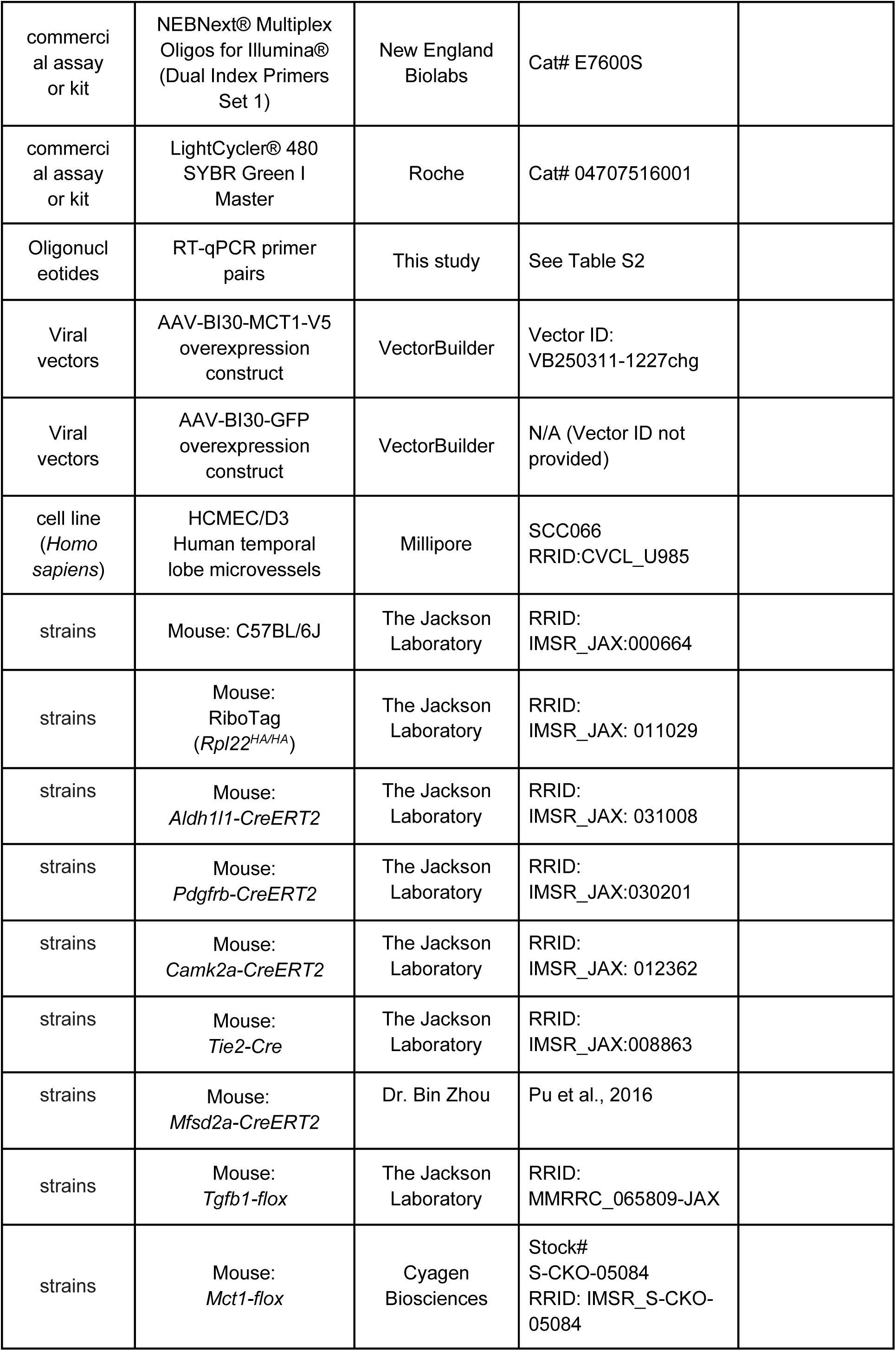
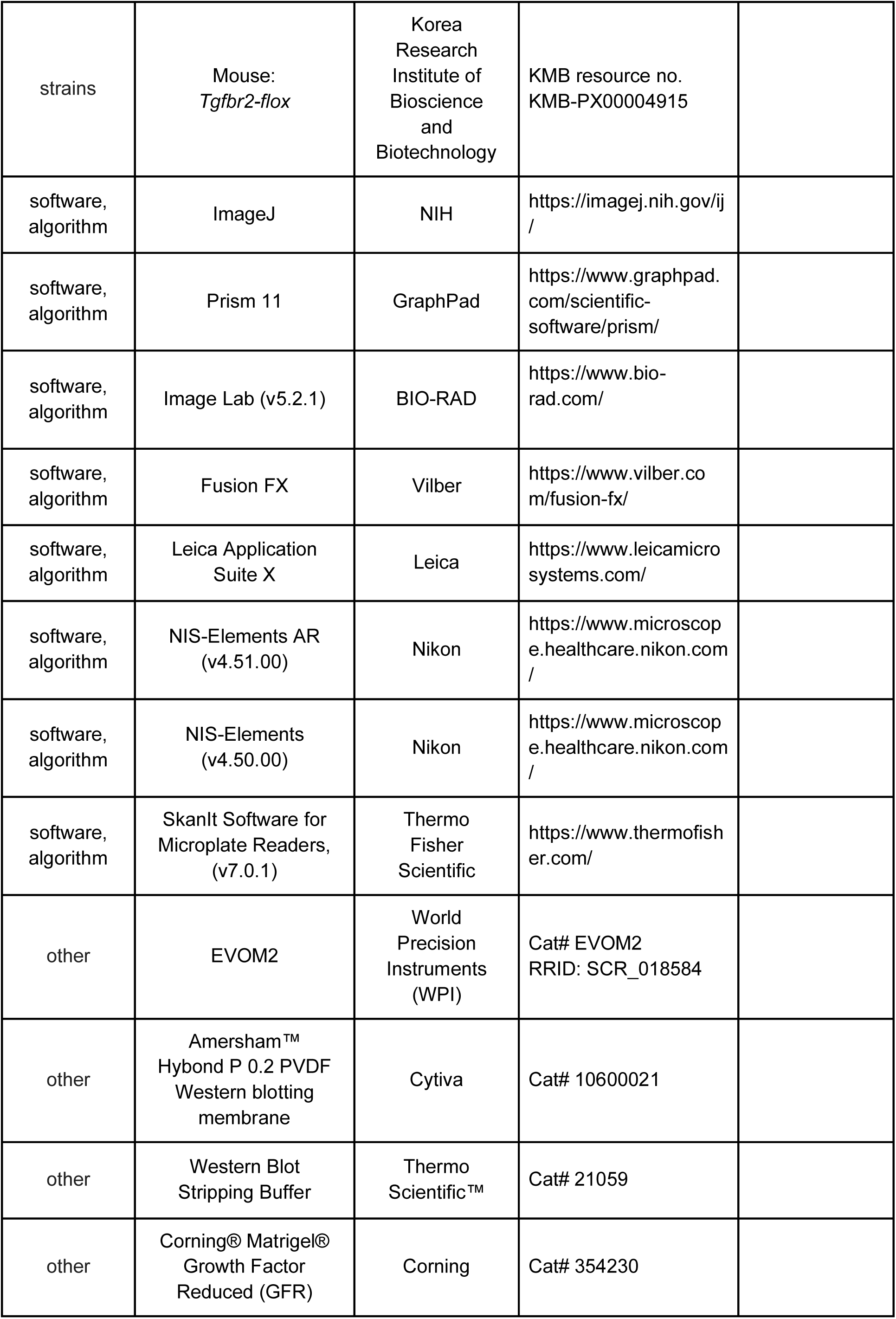
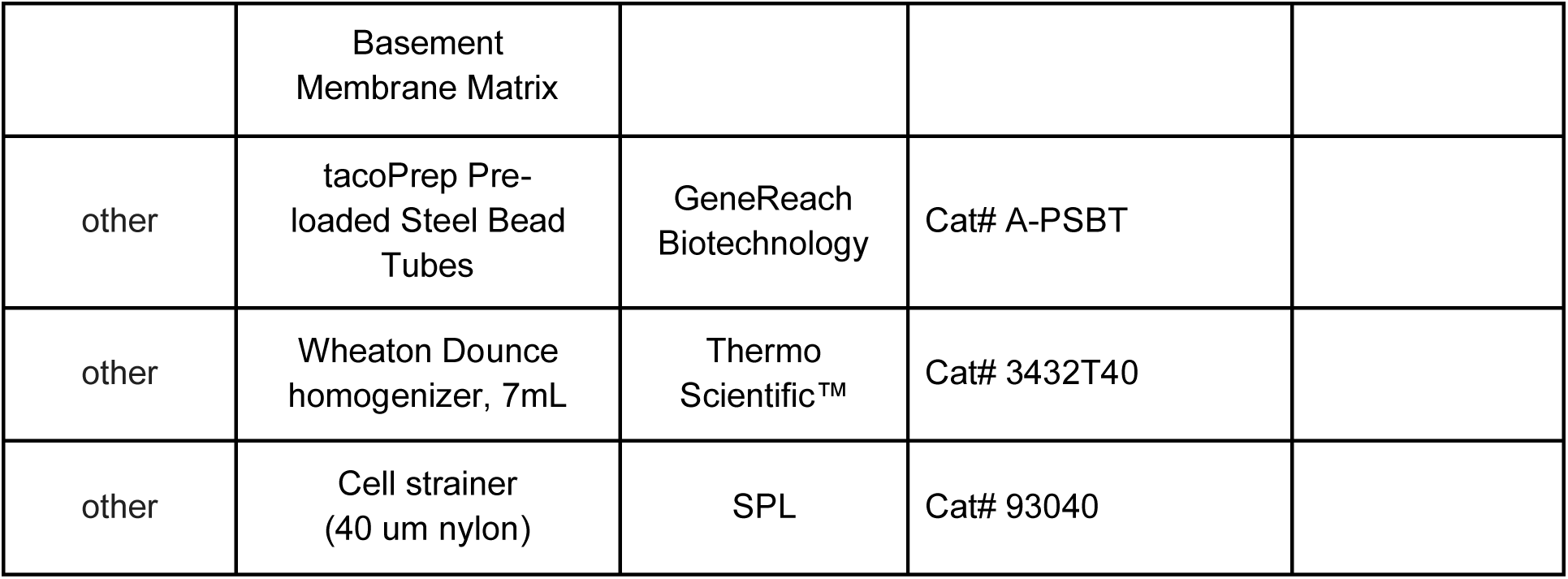

