## Supplementary_Information for "Visual experience-driven switch in endothelial metabolic transporters is essential for cerebrovascular maturation"

### Supplementary Figure 1

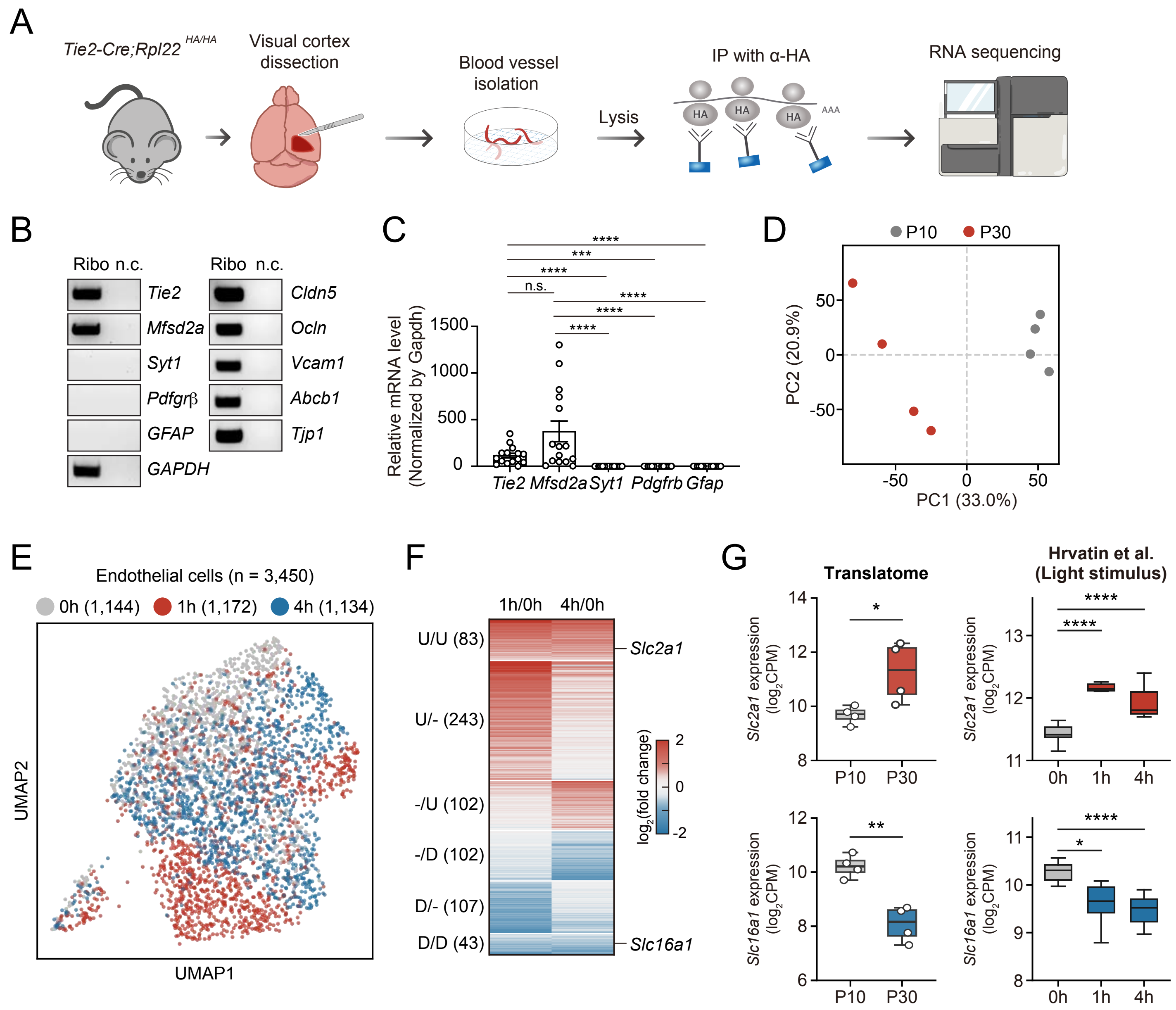


Related to Figure 1.

### Supplementary Figure 2

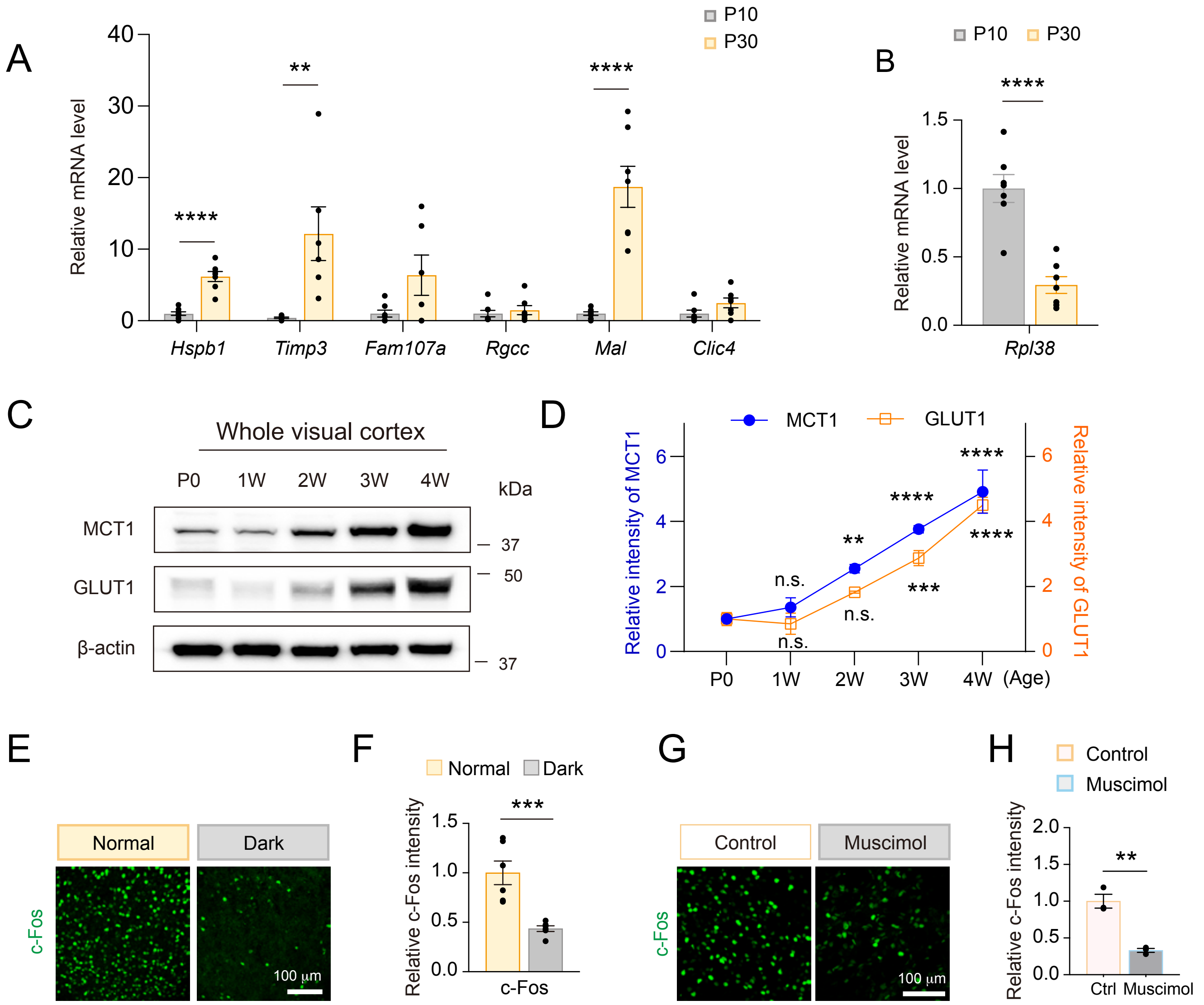


Related to Figure 1.

### Supplementary Figure 3

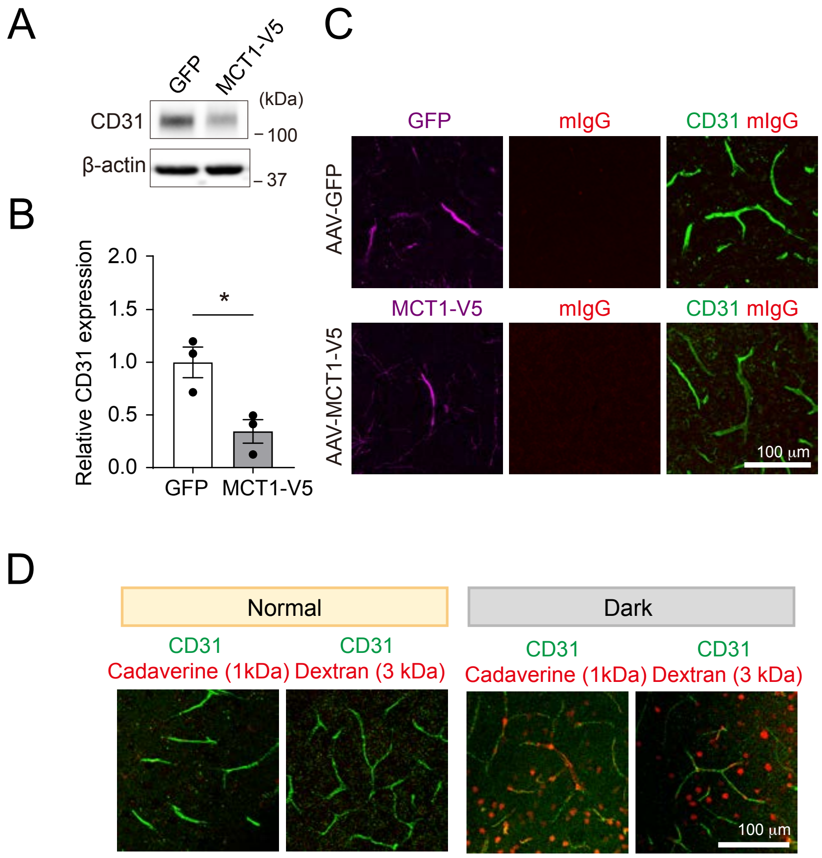


Related to Figure 2.

### Supplementary Figure 4

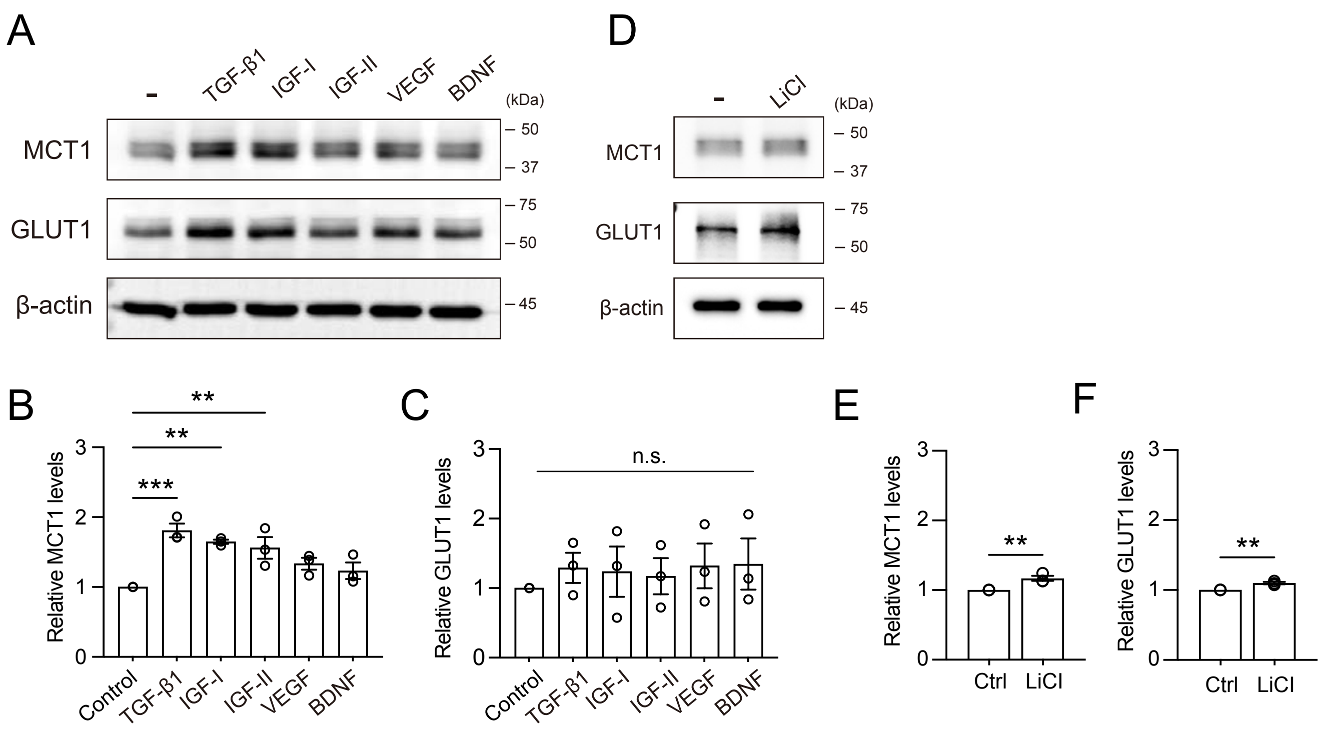


Related to Figure 3.

### Supplementary Figure 5

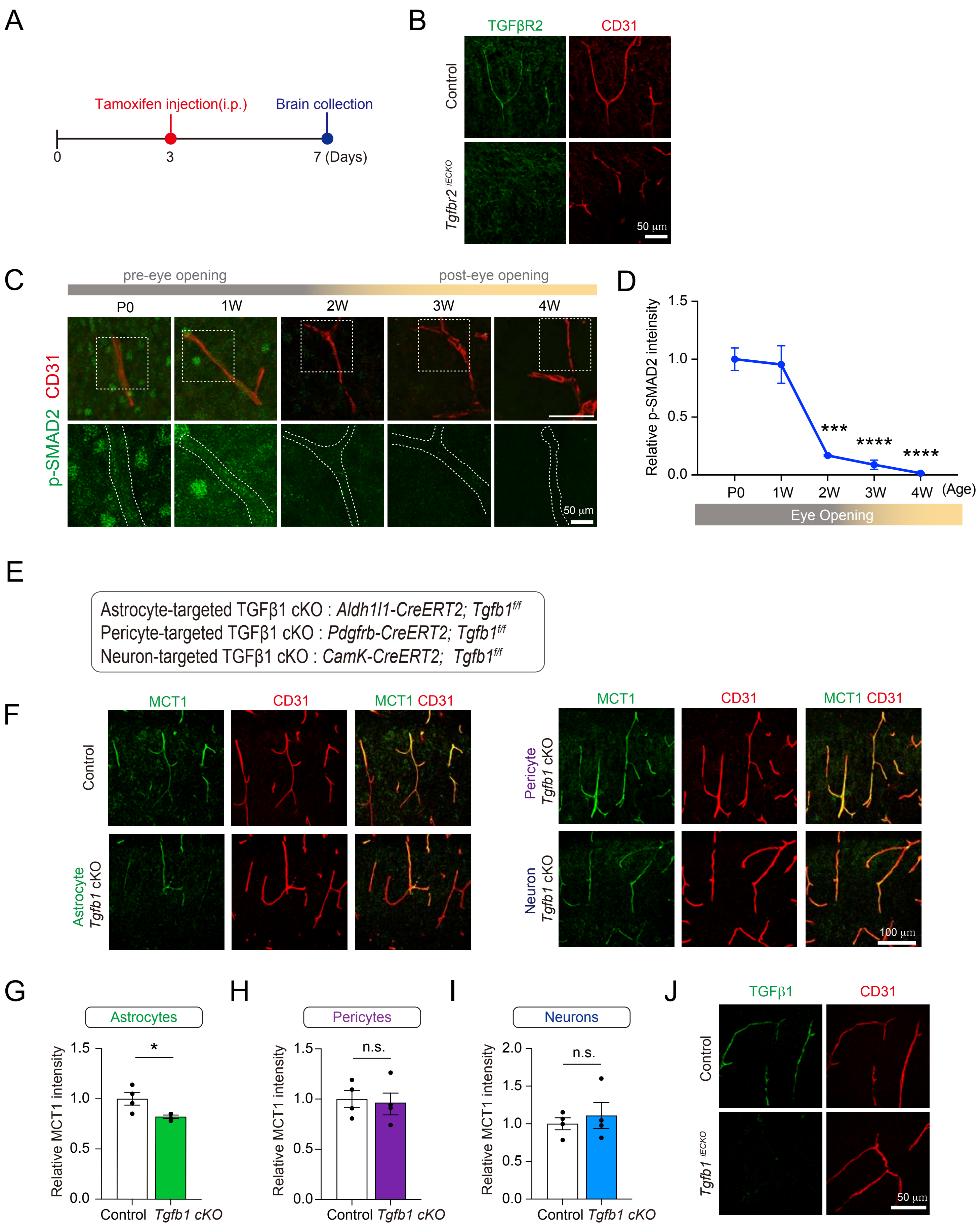


Related to Figure 3.

### Supplementary Figure 6

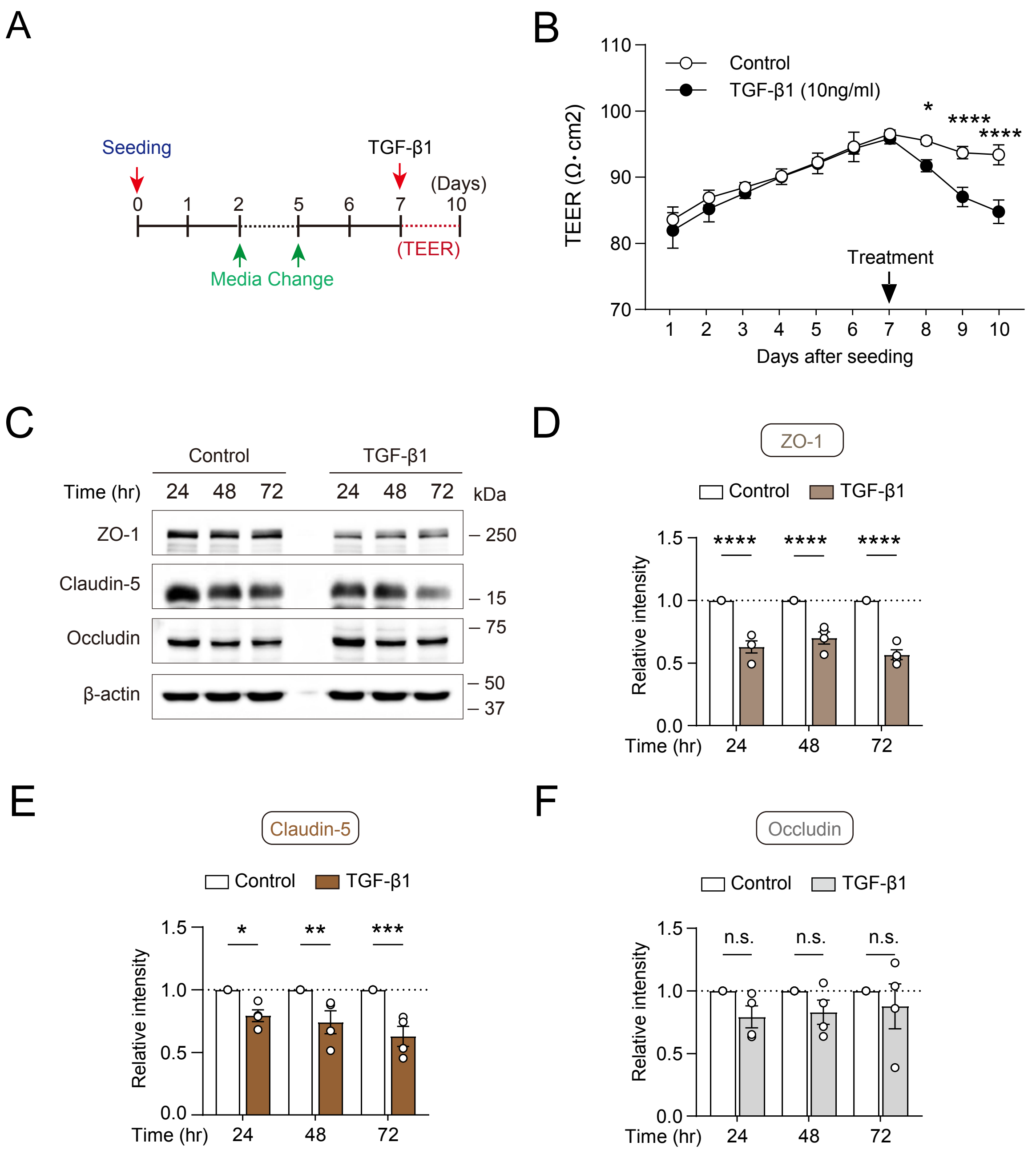


Related to Figures 2 and 3.

### Supplementary Figure 7

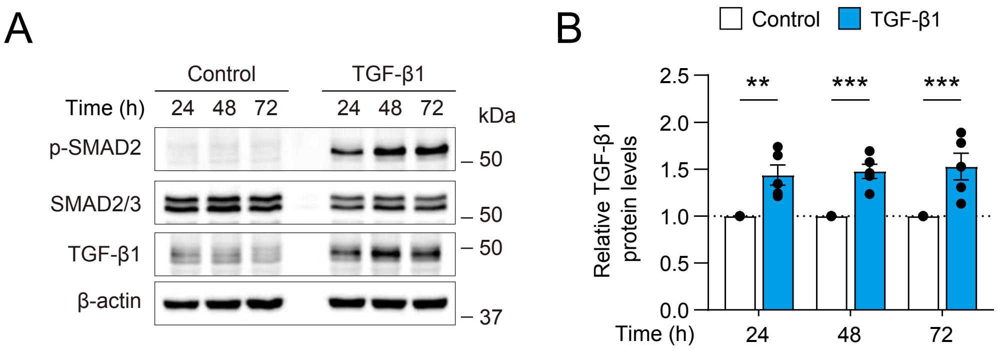


Related to Figure 4.

### Supplementary Figure 8

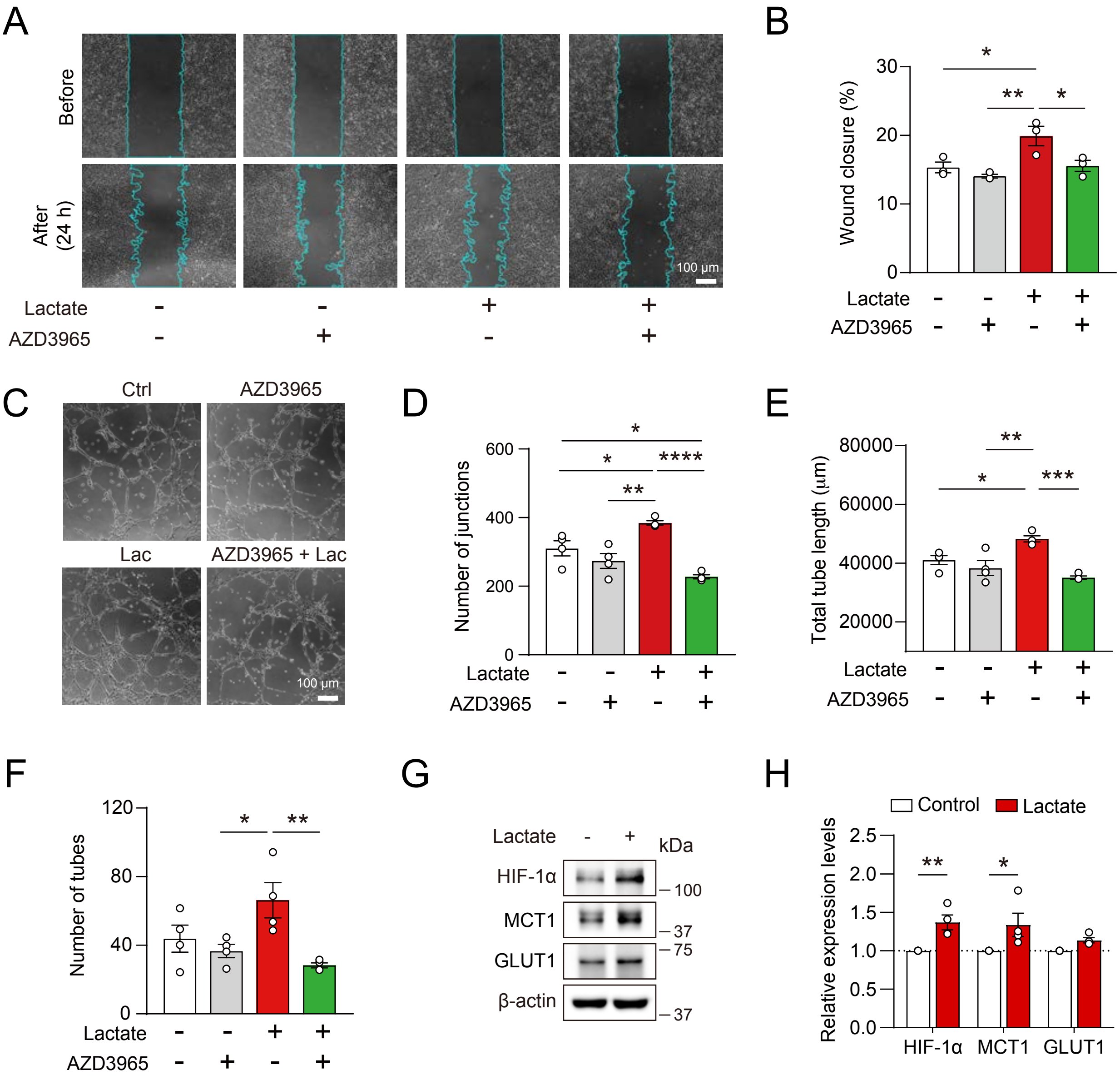


Related to Figure 5.

### Supplementary Figure legends

**Supplementary Figure 1. Gene expression profiling of brain endothelial cells in response to visual experience**

(A) Schematic of RiboTag-based endothelial translatome profiling of the visual cortex from *Tie2-Cre; $Rpl22^{HA/HA}$* mice, performed as previously described[25].

(B) Representative RT-PCR analysis of endothelial and non-endothelial cell-type markers using RNA isolated from the visual cortices by the RiboTag method. *Tie2, Mfsd2a, Cldn5, Ocln, Vcam1, Abcb1,* and *Tjp1* were used as endothelial cell markers, *Syt1* as a neuronal marker, *Pdgfrb* as a pericyte marker, and *Gfap* as an astrocyte marker. n.c., negative control.

(C) RT-qPCR analysis of cell-type-specific marker genes to assess the endothelial specificity of RNA isolated by the RiboTag method. *Tie2* and *Mfsd2a* were used as endothelial cell markers, *Syt1* as a neuronal marker, *Pdgfrb* as a pericyte marker, and *Gfap* as an astrocyte marker. n = 15 biological replicates.

(D) Principal component analysis of brain endothelial translatomes from the visual cortex at P10 (grey) and P30 (red). The proportion of variance explained by each component is indicated on the axes. n = 4 mice per group.

(E) Uniform manifold approximation and projection (UMAP) of endothelial cells extracted from visual cortex single-cell RNA sequencing data of mice exposed to light for 0, 1 or 4 h[27], colored by stimulation condition. The number of cells in each condition is indicated in parentheses.

(F) Heat map of light-responsive genes in brain endothelial cells, grouped by their expression pattern at 1 h and 4 h relative to 0 h (U, upregulated; D, downregulated; –, unchanged). The number of genes in each group is indicated in parentheses. Slc2a1 and Slc16a1 are highlighted.

(G) Expression of Slc2a1 (top) and Slc16a1 (bottom) in the endothelial translatome (left; P10 and P30, n = 4 mice per group) and in endothelial cells from single-cell RNA sequencing (right: n = 9 at 0 h and n = 7 each at 1 h and 4 h). Expression values in the single-cell dataset are summed across all endothelial cells of each mouse. Box plots show the median and interquartile range, with whiskers extending to 1.5 times the interquartile range.

Data are presented as mean ± SEM; *$p < 0.05$, **$p < 0.01$, ***$p < 0.001$, ****$p < 0.0001$ and n.s., not significant. Related to Figure 1.

**Supplementary Figure 2. Validation of top light-responsive targets identified by BEC translatome analysis and confirmation of neural activation**

(A and B) RT-qPCR analysis of selected candidate genes using endothelial RNA isolated from P10 and P30 visual cortices by the RiboTag method. n = 6–8 samples per group.

(C) Representative western blots of MCT1 and GLUT1 in whole visual cortex lysates at P0, 1W, 2W, 3W, and 4W. β-actin was used as a loading control.

(D) Quantification of MCT1 and GLUT1 protein levels at the indicated postnatal ages, normalized to β-actin. n = 3 mice per age group. One-way ANOVA followed by Dunnett's multiple-comparisons test.

(E) Representative immunofluorescence images of c-Fos in the visual cortex of normally reared and dark-reared mice. Scale bar, 100 μm.

(F) Quantification of relative c-Fos fluorescence intensity in the visual cortex of normally reared and dark-reared mice. n = 4 mice per group, with four sections analyzed per mouse. Unpaired two-tailed Student's *t* test.

(G) Representative immunofluorescence images of c-Fos in the visual cortex of control and muscimol-treated mice. Scale bar, 100 μm.

(H) Quantification of relative c-Fos fluorescence intensity in the visual cortex of control and muscimol-treated mice. n = 3 mice per group. Unpaired two-tailed Student's *t* test.

Data are presented as mean ± SEM; $^{**}p < 0.01$, $^{***}p < 0.001$, $^{****}p < 0.0001$ and n.s., not significant. Related to Figure 1.

**Supplementary Figure 3. Visual deprivation or sustained endothelial MCT1 expression impairs vascular maturation**

(A) Representative western blot of CD31 in whole brain lysates from AAV-GFP- and AAV-MCT1-V5-injected mice. β-actin was used as a loading control.

(B) Quantification of relative CD31 protein levels, normalized to β-actin. n = 3 mice per group. Unpaired two-tailed Student's *t* test.

(C) Representative immunofluorescence images of endogenous mouse IgG (mIgG) in the cerebral cortex of AAV-GFP- and AAV-MCT1-V5-injected mice. Similar findings were observed in three mice per group. Scale bar, 100 μm.

(D) Representative images of BBB permeability assessed using cadaverine (1 kDa) and dextran (3 kDa) in the visual cortex of normally reared and dark-reared mice. Similar findings were observed in three mice per group. Scale bar, 100 μm.

Data are presented as mean ± SEM; *$p < 0.05$. Related to Figure 2.

**Supplementary Figure 4. Screening of signaling factors regulating MCT1 and GLUT1 expression in brain endothelial cells**

(A) Representative western blots of MCT1 and GLUT1 in hCMEC/D3 cells treated with TGF-β1 (10 ng/mL), IGF-I (100 ng/mL), IGF-II (100 ng/mL), VEGF (100 ng/mL), or BDNF (100 ng/mL) for 72 h. β-actin was used as a loading control.

(B and C) Quantification of MCT1 (B) and GLUT1 (C) protein levels normalized to β-actin in hCMEC/D3 cells treated with the indicated factors. n = 3 biological replicates per group. One-way ANOVA followed by Dunnett's multiple comparisons test.

(D) Representative western blots of MCT1 and GLUT1 in hCMEC/D3 cells treated with lithium chloride (LiCl, 20 mM) for 72 h. β-actin was used as a loading control.

(E and F) Quantification of MCT1 (E) and GLUT1 (F) protein levels normalized to β-actin in control and LiCl-treated cells. n = 3 biological replicates per group. Unpaired two-tailed Student's *t* test.

Data are presented as mean ± SEM; **$p < 0.01$, ***$p < 0.001$, and n.s., not significant. Related to Figure 3.

**Supplementary Figure 5. Developmental TGF-β signaling and endothelial-derived TGF-β1 regulate MCT1 expression in brain endothelial cells**

(A) Experimental scheme for inducible endothelial-specific deletion of *Tgfbr2*. Tamoxifen was administered intraperitoneally at P3, and brains were collected at P7.

(B) Representative immunofluorescence images of TGFBR2 and CD31 in the visual cortex of control and endothelial-specific *Tgfbr2* conditional knockout (*Tgfbr2* $^{\text{iECKO}}$) mice to confirm endothelial TGFBR2 deletion. Similar findings were observed in two mice per group. Scale bar, 50 μm.

(C) Representative immunofluorescence images of phosphorylated SMAD2 (p-SMAD2) and CD31 in the visual cortex at the indicated postnatal ages. Boxed regions are shown at higher magnification, and vessel boundaries are indicated by dashed outlines. The approximate timing of eye opening is indicated above the images. Scale bar, 50 μm.

(D) Quantification of relative endothelial p-SMAD2 fluorescence intensity in blood vessels of the visual cortex at the indicated postnatal ages. The approximate timing of eye opening is indicated below the graph. n = 3 mice per age, with two sections analyzed per mouse. One-way ANOVA with Tukey's multiple-comparisons test.

(E) Genetic strategies for inducible cell-type-specific deletion of *Tgfb1* in astrocytes, pericytes, and neurons using *Aldh1l1-CreERT2*; *Tgfb1*$^{fl/fl}$, *Pdgfrb-CreERT2*; *Tgfb1*$^{fl/fl}$, and *Camk2a-CreERT2*; *Tgfb1*$^{fl/fl}$ mice, respectively.

(F) Representative immunofluorescence images of MCT1 and CD31 in the visual cortex of control mice and mice with astrocyte-, pericyte-, or neuron-specific conditional deletion of *Tgfb1*. Scale bar, 100 μm.

(G–I) Quantification of relative endothelial MCT1 fluorescence intensity in the visual cortex of astrocyte-specific (G), pericyte-specific (H), and neuron-specific (I) *Tgfb1* conditional knockout mice and their respective controls. n = 4 mice per group, with two sections analyzed per mouse. Unpaired two-tailed Student's *t* test.

(J) Representative immunofluorescence images of TGF-β1 and CD31 in the visual cortex of control and endothelial-specific *Tgfb1* conditional knockout (*Tgfb1* $^{iECKO}$) mice. Scale bar, 50 μm.

Data are presented as mean ± SEM; $*p < 0.05$, $***p < 0.001$, $****p < 0.0001$ and n.s., not significant. Related to Figure 3.

**Supplementary Figure 6. Canonical TGF-β/Smad signaling compromises BBB integrity in cultured brain endothelial cells**

(A) Experimental scheme for assessment of endothelial barrier integrity by transendothelial electrical resistance (TEER). hCMEC/D3 cells were seeded at day 0, and culture medium was changed at the indicated time points. TGF-β1 treatment was initiated on day 7, followed by TEER measurements at the indicated time points.

(B) TEER measurements of hCMEC/D3 monolayers following control or TGF-β1 treatment at the indicated time points. n = 3 biological replicates per group. Two-way ANOVA followed by Sidak's multiple-comparisons test.

(C) Representative western blot analysis of ZO-1, Claudin-5, and Occludin in hCMEC/D3 cells under the same experimental conditions as the TEER assay following control or TGF-β1 treatment for 24, 48, or 72 h. β-actin was used as a loading control.

(D–F) Quantification of ZO-1 (D), Claudin-5 (E), and Occludin (F) protein levels normalized to β-actin in control and TGF-β1-treated hCMEC/D3 cells at 24, 48, and 72 h. n = 4 biological replicates per group. Two-way ANOVA followed by Sidak's multiple-comparisons test.

Data are presented as mean ± SEM; $^{*}p < 0.05$, $^{**}p < 0.01$, $^{***}p < 0.001$, $^{****}p < 0.0001$ and n.s., not significant. Related to Figures 2 and 3.

**Supplementary Figure 7. TGF-β signaling induces its own ligand expression in brain endothelial cells**

(A) Representative western blots of phosphorylated SMAD2 (p-SMAD2), total SMAD2/3, and TGF-β1 in hCMEC/D3 cells treated with TGF-β1 for 24, 48, or 72 h. β-actin was used as a loading control. Note that TGF-β1 treatment significantly increases p-SMAD2 levels.

(B) Quantification of TGF-β1 protein levels in control and TGF-β1-treated cells at the indicated time points, normalized to β-actin. n = 5 biological samples per group. Two-way ANOVA followed by Sidak's multiple-comparisons test.

Data are presented as mean ± SEM; $^{**}p < 0.01$ and $^{***}p < 0.001$. Related to Figure 4.

**Supplementary Figure 8. Lactate-MCT1 axis promotes vascular growth in cultured brain endothelial cells**

(A) Representative images of scratch-wound migration assays in hCMEC/D3 cells treated with L-lactate (10 mM), the MCT1 inhibitor AZD3965 (1 μM), or both. Scale bar, 100 μm.

(B) Quantification of endothelial cell migration under the indicated treatment conditions. n = 3 biological replicates per group. One-way ANOVA followed by Tukey's multiple comparisons test.

(C) Representative images of endothelial tube formation following treatment with lactate, AZD3965, or both. Scale bar, 100 μm.

(D–F) Quantification of the number of junctions (D), total tube length (E), and number of tubes (F) per field under the indicated treatment conditions. n = 3 biological replicates per group. One-way ANOVA followed by Tukey's multiple comparisons test.

(G) Representative western blots of HIF-1α, MCT1, and GLUT1 in hCMEC/D3 cells with or without L-lactate treatment. β-actin was used as a loading control.

(H) Densitometric quantification of HIF-1α, MCT1, and GLUT1 protein levels normalized to β-actin. n = 4 biological samples per group. Two-way ANOVA followed by Dunnett's multiple-comparisons test. Data are presented as mean ± SEM; *$p < 0.05$, **$p < 0.01$, ***$p < 0.001$, ****$p < 0.0001$ and n.s., not significant. Related to Figure 5.
